# Structure of the two-component S-layer of the archaeon *Sulfolobus acidocaldarius*

**DOI:** 10.1101/2022.10.07.511299

**Authors:** Lavinia Gambelli, Mathew McLaren, Rebecca Conners, Kelly Sanders, Matthew C. Gaines, Lewis Clark, Vicki Gold, Daniel Kattnig, Mateusz Sikora, Cyril Hanus, Michail N. Isupov, Bertram Daum

## Abstract

Surface layers (S-layers) are resilient two-dimensional protein lattices that encapsulate many bacteria and most archaea. In archaea, S-layers usually form the only structural component of the cell wall and thus act as the final frontier between the cell and its environment. Therefore, S-layers are crucial for supporting microbial life. Notwithstanding their importance, little is known about archaeal S-layers at the atomic level. Here, we combined single particle cryo electron microscopy (cryoEM), cryo electron tomography (cryoET) and Alphafold2 predictions to generate an atomic model of the two-component S-layer of *Sulfolobus acidocaldarius*. The outer component of this S-layer (SlaA) is a flexible, highly glycosylated, and stable protein. Together with the inner and membrane-bound component (SlaB), they assemble into a porous and interwoven lattice. We hypothesize that jackknife-like conformational changes, as well as pH-induced alterations in the surface charge of SlaA, play important roles in S-layer assembly.

## Introduction

The prokaryotic cell envelope includes a cytoplasmic membrane and a cell wall, which provide the structural integrity to the cell and mediate the interaction between the extracellular and intracellular environment. The cell wall differs in composition and structure across prokaryotes^1, 2^. In bacteria, a peptidoglycan (murein) layer encapsulates the cytoplasmic membrane, and this is in turn enclosed by a second membrane in Gram-negative bacteria^1^. Generally, the archaeal cell wall lacks an outer membrane, but a variety of cell wall elements, including pseudomurein, methanochondroitin and protein sheaths have been described^3^. Most prokaryotes exhibit a porous glycoproteic surface layer (S-layer) as the outermost component of their cell wall. In archaea, S-layers are the simplest and most commonly found cell wall structure^2–4^.

The prokaryotic cell envelope is exposed to a variety of environmental conditions, which, in the case of extremophiles, can be unforgiving (low/high pH, high temperature and salinity). Therefore, S-layers reflect the cellular need for both structural and functional plasticity, allowing archaea to thrive in diverse ecosystems. Archaeal S-layers maintain the cell shape under mechanical, osmotic and thermal stress, selectively allow molecules to enter or leave the cell, and create a quasiperiplasmic compartment (similar to the periplasmic space in Gram-negative bacteria)^2–4^. S-layer glycoproteins are also involved in cell-cell recognition^5^ and mediate virus-host interactions^6^.

Structurally, an S-layer is a pseudocrystalline array of (glyco)proteins (surface layer proteins, SLPs). The ordered nature of an S-layer is what sets it apart from other protein sheaths^3^. S-layers usually consist of thousands of copies of one SLP species. These SLPs self-assemble on the cell surface predominantly at mid-cell^7, 8^, giving rise to an oblique (p1), square (p2, p4) or hexagonal (p3, p6) symmetry^9^. In archaea, the hexagonal symmetry is the most common^2^. The S-layer is highly porous. Depending on the species, the pores can occupy up to about 70 % of the S-layer surface and have different sizes and shapes^2^. Such an assembly provides a highly stable and flexible 2D lattice^10, 11^. Archaeal SLPs range from 40 - 200 kDa in molecular mass and show little sequence conservation^7^. The most common post-translational modification of SLPs is glycosylation. Most archaeal SLPs are N- and/or O-glycosylated and the composition of the glycans is highly diverse^2, 4^. Thermophilic and hyperthermophilic archaea show a higher number of glycosylation sites on SLPs compared to mesophilic archaea, suggesting that glycans support thermostability^12^. Another common aspect of archaeal S-layers is their binding of divalent metal ions^11, 13, 14^, which have been shown to be essential for S-layer assembly and anchoring in bacteria^15, 16^. Atomic models on archaeal S-layers have been reported for domains of *Methanosarcina* SLPs ^17, 18^, and more recently, an atomic structure of the *Haloferax volcanii* S-layer has been described^14^.

*Sulfolobus acidocaldarius* is a hyperthermophilic and acidophilic archaeon of the Crenarchaeota phylum and thrives in acidic thermal soils and hot springs worldwide. It grows at pH ∼2-3 and temperatures ranging from 65 °C to 90 °C^19^. The *Sulfolobus* S-layer is composed of two repeating glycoproteins, SlaA and SlaB. In *S. acidocaldarius*, SlaA contains 1,424 amino acids and has a molecular mass of 151 kDa, whereas SlaB comprises 475 amino acids and has a mass of 49.5 kDa^20^. Comparative sequence analysis and molecular modelling predicted that SlaA is a soluble protein rich in β-strands^21^. On the other hand, SlaB has been predicted to contain three consecutive β-sandwich domains at the N-terminus and a membrane-bound coiled-coil domain at the C-terminus^21^. Across the Sulfolobales, SlaA shows higher sequence and structural variability compared to SlaB^21^. Early 2D crystallography and electron microscopy experiments described the *S. acidocaldarius* S-layer as a “smooth”, highly porous, hexagonal (p3) lattice^20, 22^. Recently, we investigated the architecture of the *S. acidocaldarius* S-layer by electron cryo-tomography (cryoET)^23^. The S-layer has a bipartite structure with SlaA and SlaB forming the extracellular- and intracellular-facing layers, respectively. Dimers of SlaA and trimers of SlaB assemble around hexagonal and triangular pores, creating a ∼30 nm thick canopy-like framework. However, the resolution was limited and secondary structure details were unresolved. *Sulfolobus* mutants lacking SlaA and/or SlaB show morphological aberrations, higher sensitivity to hyperosmotic stress and alterations of the chromosome copy number, suggesting that in these species the S-layer plays key roles in cell integrity, maintenance, and cell division^24^.

Here, we investigated the *S. acidocaldarius* S-layer and its components using a combination of single particle cryoEM and cryoET. We solved the atomic structure of SlaA and investigated its stability across extreme pH ranges. Moreover, we combined cryoEM data and Alphafold2 to build a complete *in situ* atomic model of this S-layer and propose insights into their dynamics and assembly.

## Results

### Structure and N-glycosylation of SlaA_30-1,069_ at acidic pH

To solve the structure of the *S. acidocaldarius* SLP SlaA, we disassembled the S-layer by changing the pH from acidic to basic and purified the native protein using size exclusion chromatography. Suspensions of the protein at pH 4 were plunge frozen using cryoEM grids. The acidic pH was chosen to account for the natural conditions in which *S. acidocaldarius* thrives. The structure of SlaA was determined from cryoEM movies, using the single particle analysis (SPA) pipeline in Relion 3.1^25^ (Supplementary Fig. 1, 2a,d; Supplementary Table 1). The final cryoEM map had a global resolution of 3.1 Å (Supplementary Fig. 3a, b; 4a).

Because SlaA has virtually no homology with other structurally characterised proteins, the cryoEM map was used to build an atomic model *de novo* (Fig. 1a; Supplementary Fig. 4b; Supplementary Movie 1). Residues 30 to 1,069 (∼ 70 % of the sequence) were clearly defined in the cryoEM density map. The N-terminal signal peptide (predicted to be residues 1-24) is cleaved prior to S-layer assembly^21^. A few N-terminal residues and residues 1,070-1,424 at the C-terminus were not resolved by SPA due to their high flexibility (Supplementary Fig. 5; Supplementary Movie 2).

**Figure 1.**
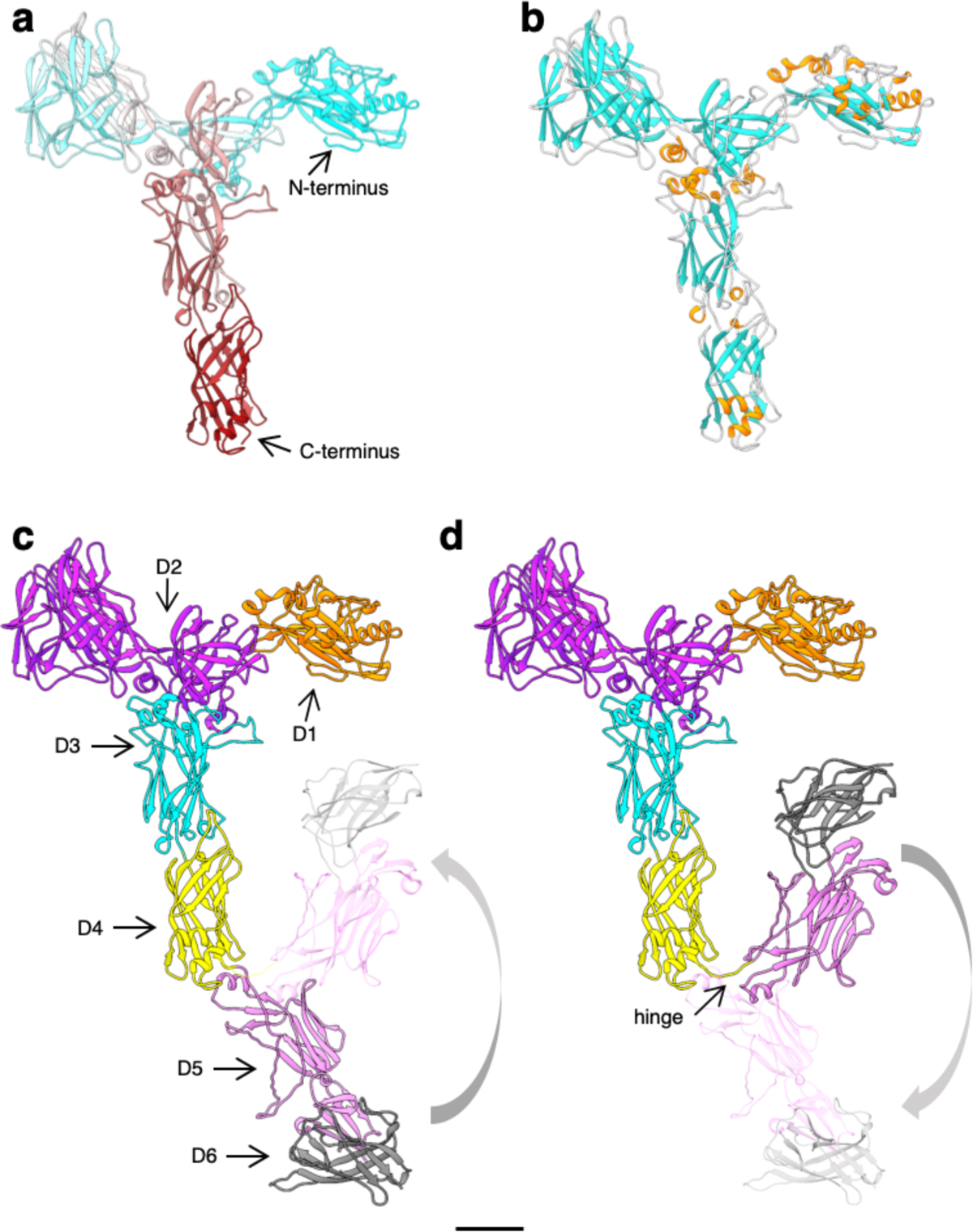
Atomic model of *S. acidocaldarius* S-layer protein SlaA at pH 4. **a**, SlaA30-1,069 atomic model in ribbon representation and cyan-grey-maroon colours (N-terminus, cyan; C-terminus, maroon). **b**, SlaA30-1,069 atomic model showing the α-helices in orange, the β-strands in cyan and the loop regions in grey. **c** and **d**, SlaA atomic models highlighting six domains: D130-234 (orange), D2235-660,701-746 (purple), D3661-700,747-914 (cyan), D4915-1,074 (yellow), D51,075-1,273 (pink), and D61,274-1,424 (grey). D5 and D6 were predicted using Alphafold. The flexibility of a link between domains D4 and D5 is highlighted showing stretched (c) and flapped (d) conformations. Scale bar, 20 Å.

SlaA_30-1,069_ is a Y-shaped protein. It consists mostly of β-strands and contains only a few short α-helices (Fig. 1b, Supplementary Fig. 4c,d). The polypeptide chain is arranged into four domains (D1_30-234_, D2_235-660,701-746_, D3_661-700,747-914_, D4_915-1,069_) <u>defined by</u> SWORD^26^ (Fig. 1c).

Of those domains, only D4 shows significant similarity to known structures – to domain 3 of complement C5 (PDB ID: 4E0S) according to DALI^27^. A disulphide bond links D3 and D4 (Cys_677_-Cys_1,017_) (Supplementary Fig. 4d), however, the density of this bond is not visible in the cryoEM map, likely due to electron beam damage^28^.

The structure of the missing C-terminus (SlaA_914-1,424_) was predicted (including D4 to aid alignment) using Alphafold^29^ and revealed two additional β-domains, D5 and D6, (Supplementary Fig. 6). Alphafold predicted five different conformations of SlaA_914-1,424_, which differed with regards to the position of D5-D6 relative to D1-D4, suggesting an in-plane flexibility between these two parts of the protein around a hinge (amino acids A_1,067_-L_1,071_) between D4 and D5 (Supplementary Fig. 6). Similar conformations were also observed in 2D classes of our cryoEM dataset (Supplementary Fig. 5), substantiating the Alphafold predictions in Supplementary Fig. 6. The extremes of the conformational space of SlaA are shown in Fig. 1c and 1d. These describe stretched (“open”) and flapped (“closed”) conformations. It is probable that this jackknife-like flexibility is a prerequisite for the protein to assemble into the S-layer.

SlaA is expected to be highly glycosylated; its sequence contains 31 predicted N-glycosylation sites^30^. Our cryoEM map of SlaA_30-1,069_ shows 19 glycan densities (Fig. 2), largely in agreement with the prediction of 20 sequons located in this portion of the protein^30^. The 19 glycosylated Asn residues in SlaA_30-1,069_ are listed in Fig. 2e. The remaining predicted glycosylation sites are located in domains D5 and D6, in which eight sites were confirmed to be glycosylated by mass spectrometry analysis^30^. Therefore, the entire SlaA protein contains a total of 27 confirmed glycans.

**Figure 2.**
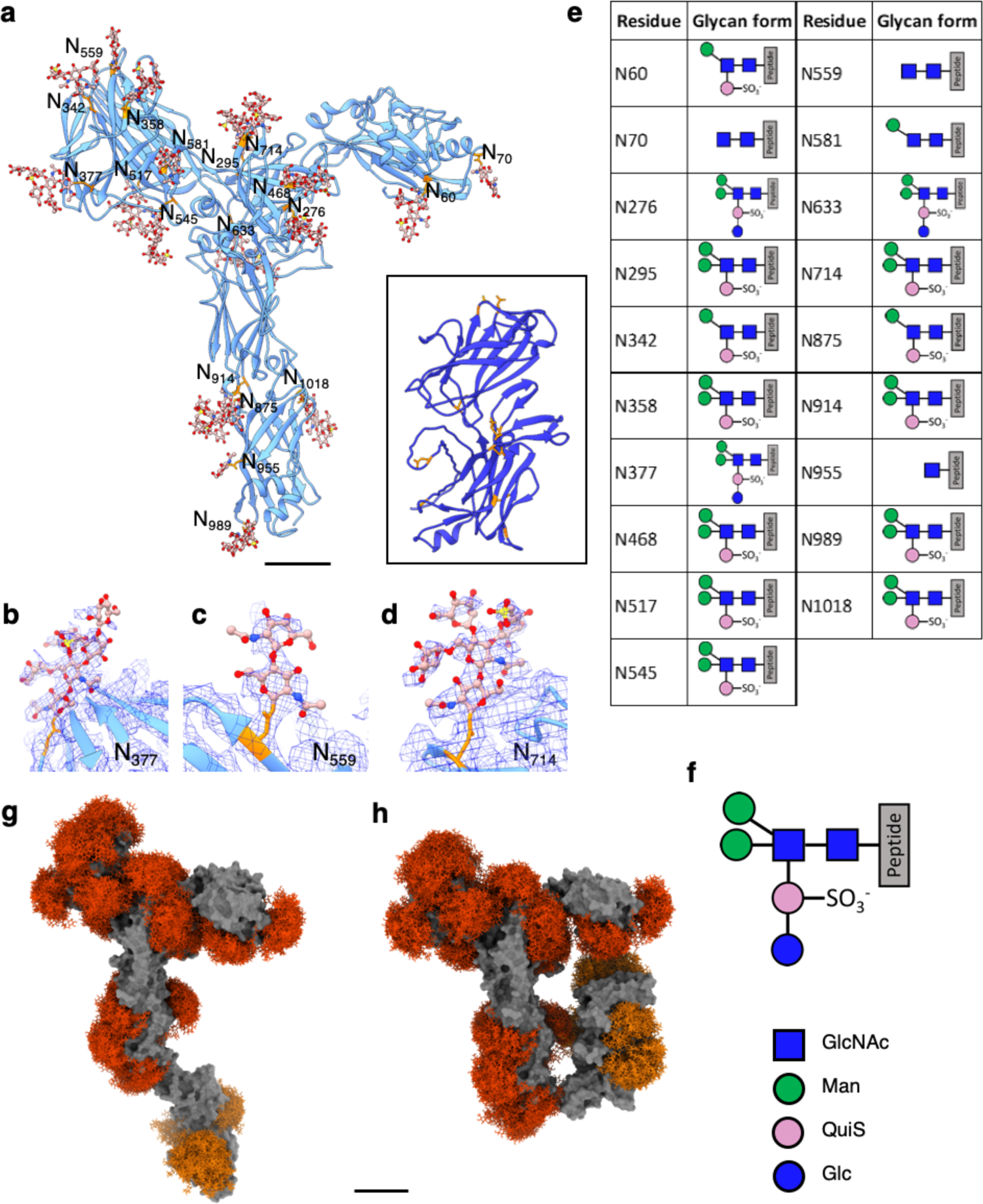
N-glycosylation of *S. acidocaldarius* SlaA. **a**, SlaA atomic model in ribbon representation. SlaA30-1,069 as solved by cryoEM is in cornflower blue; SlaA1,070-1,424 as predicted by Alphafold is in blue (boxed). All 19 Asn-bound glycan molecules (stick representation; glycans in rosy brown, Asn in orange) in SlaA30-1,069 are modelled fitting the cryoEM map. In the glycans, O atoms are shown in red, N in blue and S in yellow. Eight glycosylated Asn are highlighted (stick representation, orange) in SlaA1,070-1,424 based on Peyfoon *et al*., 2010^30^. Scale bar, 20 Å. **b**-**d** are example close-ups of glycosylation sites with superimposed cryoEM density map (blue mesh). (b) shows the full hexasaccharide on Asn377, (c) shows GlcNAc2 on Asn559, and (d) shows a pentasaccharide lacking Glc1 on Asn714. **e**, list of glycosylation sites and associated glycans of SlaA30-1,069. The schematic glycan representation (**f**) is equivalent to Peyfoon *et al*., 2010^30^. Blue square, N-acetylglucosamine; green circle, mannose; pink circle, 6-sulfoquinovose; blue circle, glucose. **g** and **h**, GlycoSHIELD models (red, orange) showing the glycan coverage of the protein (solid grey). Glycan shields corresponding to glycosylation sites visualised by cryoEM are highlighted in red, glycan shields with the Alphafold model of the SlaA C-terminus are shown in orange.

The N-glycans were modelled into the cryoEM densities based on their known chemical structure^31^. The complete glycan is a tribranched hexasaccharide, containing a 6-sulfoquinovose (QuiS). Not all glycosylation sites had clear density to model the entire hexasaccharide. Instead, several forms of apparently truncated glycans were fitted into the cryoEM map (Fig. 2b-d). Most glycans (47 %) were built as pentasaccharides, lacking the glucose bound to QuiS in the mature glycan; 15 % of the glycan pool could be modelled with the whole hexasaccharide structure.

As shown for other glycoproteins, in particular the spike proteins of coronavirus^32^, glycans are usually much more dynamic than polypeptides and rapidly explore large conformational spaces, generating potentially bulky glycan shields over hundreds of nanoseconds. To evaluate the morphology and span of such shields, a reductionist molecular dynamics simulation approach (GlycoSHIELD)^33^ was used to graft plausible arrays of glycan conformers onto open and closed conformations of SlaA monomers with D5 and D6 domains (Fig. 2 g,h). Glycan volume occupancy was comparable on the two conformations of the monomers (Fig. 2 g,h). The closed conformation enabled a slightly higher number of possible glycan conformers (Supplementary Fig. 8). Although it is not yet clear how this would affect S-layer assembly, it suggests that the glycans do not prevent the flapping of the protein (Fig. 4).

### SlaA at different pH conditions

SlaA assembly and disassembly are pH-sensitive processes^23^. A pH shift from acidic (∼pH 4) to alkaline (∼pH 10) induces the disassembly of the lattice into its component subunits. To shed light on the mechanism of S-layer assembly and disassembly, we investigated the structure of SlaA at different pH conditions. Purified SlaA proteins were frozen at pH 7 and pH 10 and their structure was determined using the SPA pipeline in Relion 3^34^ (Supplementary Fig. 9a, b; Supplementary Table 1) and 3.1 (Supplementary Fig. 10; Supplementary Table 1; Supplementary figure 2b-f). The resulting cryoEM maps had global resolutions of 3.9 Å for SlaA at pH 7 and 3.2 Å for SlaA at pH 10 (Fig. 3a; Supplementary Fig. 3). As for SlaA at pH 4, domains D5 and D6 were too flexible to be resolved in the cryoEM maps. Strikingly, the cryoEM maps of SlaA_30-1,069_ at the three pH conditions were virtually identical, demonstrating a remarkable pH stability of this protein. The mean r.m.s.d. value of Ca between the pH 4 and pH10 structures was 0.79 Å (min. = 0.02 Å; max. = 2.6 Å) (Fig. 3b; Supplementary Movie 3), confirming that SlaA_30-1,069_ maintains its structure unchanged across a surprisingly broad pH range. This suggests that a pH-induced conformational change in SlaA_30-1,069_ is not the cause for S-layer disassembly. However, because D5 and D6 were not resolved in our map, a structural rearrangement affecting these domains remains a possibility.

**Figure 3.**
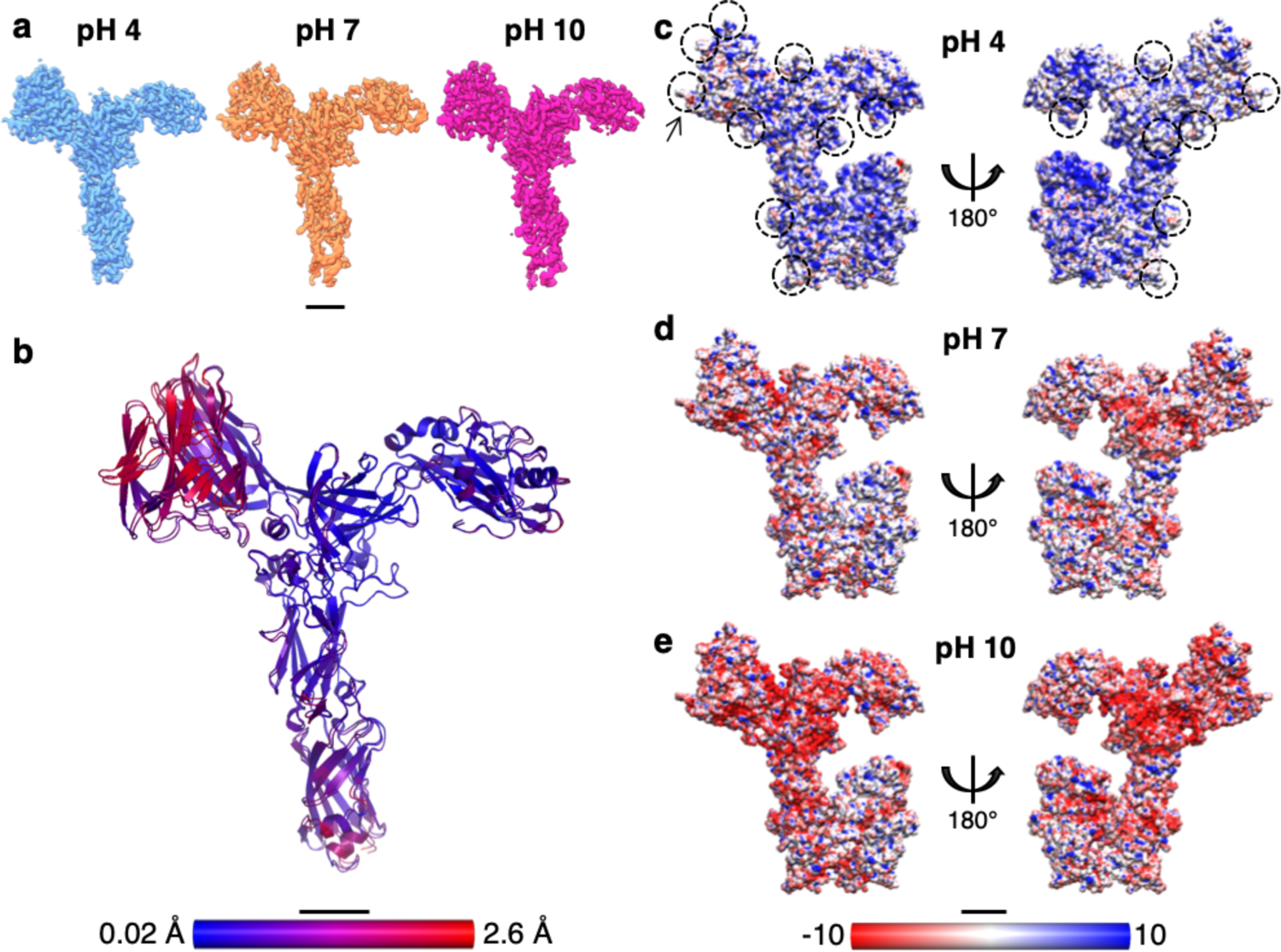
Structural comparison and electrostatic surface potentials of *S. acidocaldarius* SlaA at different pH conditions. **a**, SlaA30-1,069 cryoEM maps at pH 4 (light blue, res. 3.1 Å), pH 7 (orange, res. 3.9 Å) and pH 10 (magenta, res. 3.2 Å). **b**, r.m.s.d. (root-mean-square deviation) alignment between SlaA30-1069 atomic models at pH 4 and pH 10. Smaller deviations are shown in blue and larger deviations in red, with mean r.m.s.d. = 0.79 Å. **c-e**, electrostatic surface potentials of SlaA at pH 4 (c), pH 7 (d) and pH 10 (e). Models include Alphafold-predicted C-terminal domains (in closed conformation). Surfaces are coloured in red and blue for negatively and positively charged residues respectively. White areas represent neutral residues. In (c) some areas occupied by glycans are circled; the arrow points at one of the 6-sulfoquinovose residues displaying a negative charge at pH 4. Scale bar, 20 Å.

A variation in pH can dramatically affect protein-protein interactions by changing the overall electrostatic surface potential of the protein complex^35, 36^. An analysis of the surface charges of SlaA, including the glycans, at pH 4, 7 and 10 revealed that the overall protein charge shifts significantly from positive at pH 4 to negative at pH 10 (Fig. 3c-e). A comparison of the surface charge between glycosylated and non-glycosylated SlaA (Supplementary Fig. 11) showed that the glycans contribute considerably to the negative charge of the protein at higher pH values. This change in electrostatic surface potential may therefore be a key factor in disrupting protein-protein interactions within the S-layer, causing its disassembly.

### Atomic structure of the *S. acidocaldarius* S-layer

In a previous study, we determined the location of SlaA and SlaB within the S-layer lattice by cryoET of whole cells and isolated S-layers^23^. However, due to the limited resolution of the cryoEM maps and the lack of SlaA and SlaB atomic models, the details of the S-layer structure could not be explored. To improve on the available cryoEM map of the *S. acidocaldarius* S-layer and investigate the atomic structure of the lattice, we performed cryoET and subtomogram averaging (STA) on *S. acidocaldarius* exosomes with improved imaging conditions and processing techniques. Exosomes are naturally secreted S-layer-encapsulated vesicles, with a diameter of about 90-230 nm^37^. To analyse the *in situ* structure of the S-layer, we performed STA using Relion 3 and 3.1, followed by emClarity^38^ (Supplementary Table 1) and obtained a cryoEM map at 8.9 Å resolution (Supplementary Fig. 12a-c and 13). Finally, our structure of SlaA was fitted into the S-layer map, providing an atomic model of the assembled lattice (Fig. 4a,b; Supplementary Fig. 12 d-f).

When observed in the direction parallel to the S-layer plane, the exosome-encapsulating S-layer displays a positive curvature, with a curvature radius of the cryoEM average map of 84 nm (Fig. 4c,d). SlaA assembles into a sheet with a thickness of 95 Å. Within the long axes of the SlaA, subunits adopt an angle of about 28° with respect to the membrane plane (Fig. 4d). As a result of this inclination, effectively two zones in the SlaA assembly can be distinguished: an outer zone constituted by D1, D2, D3 and D4, and one inner zone formed by D5 and D6 (Fig. 4c,e).

**Figure 4.**
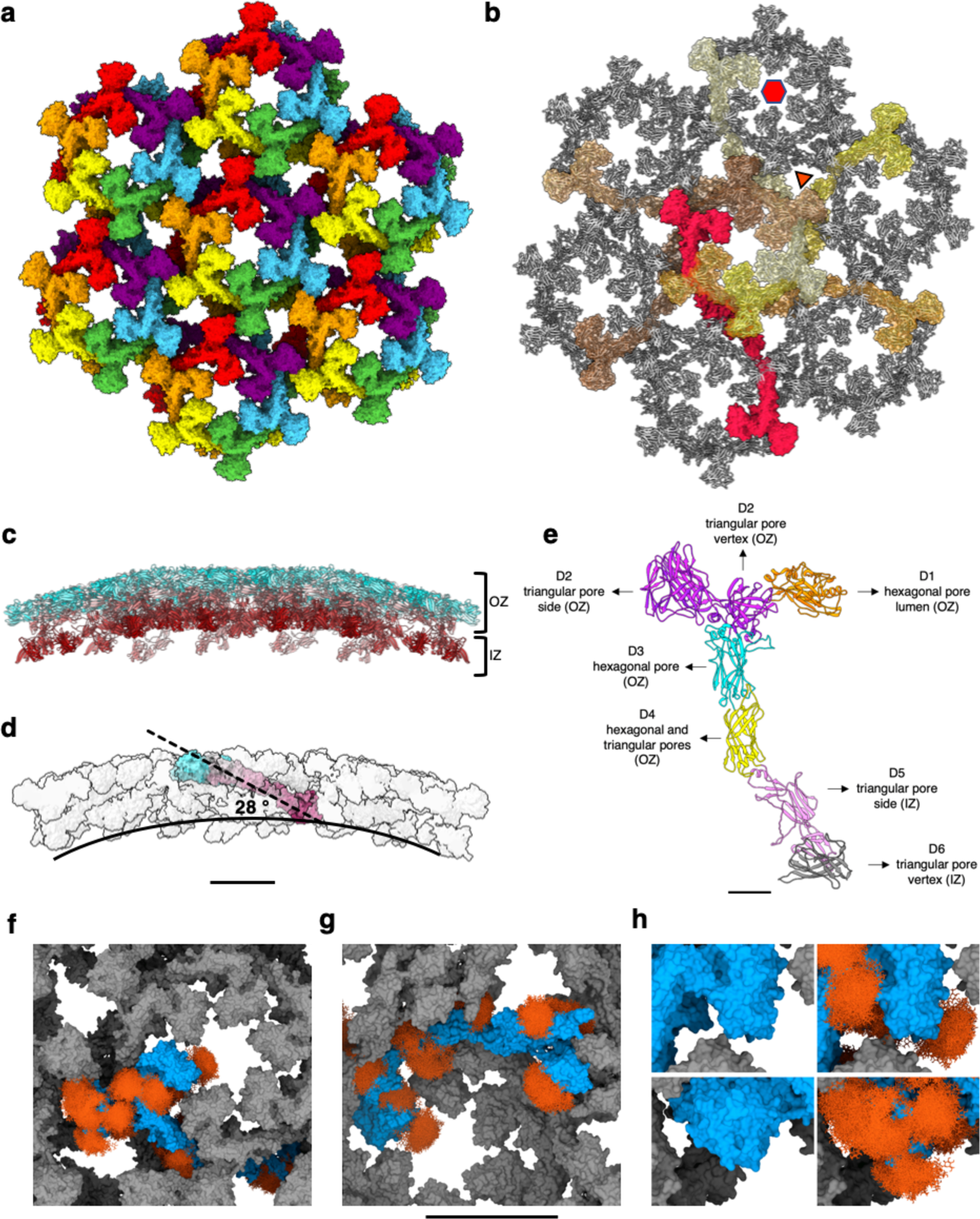
*S. acidocaldarius* SlaA assembly on the surface of exosomes. **a**, extracellular view of assembled SlaA monomers in surface representation and in randomly assigned colours. **b**, extracellular view of assembled SlaA in ribbon representation with SlaA dimers around a hexagonal pore highlighted in shades of red and surface representation. Each dimer spans two adjacent hexagonal pores. **c**, side view of the SlaA lattice (blue, N-terminus; red, C-terminus). It is possible to distinguish an outer zone (OZ) formed by domain D1, D2, D3 and D4, and an inner zone (IZ) constituted by domains D5 and D6. **d**, one SlaA monomer (surface representation, N-terminus cyan, grey, C-terminus maroon) is highlighted within assembled SlaA in surface representation. SlaA assembles with its long axis (dashed line) forming a 28° angle to the membrane plane (solid line). **e**, location of SlaA domains in assembled S-layer. The SlaA domains are highlighted in different colours: D130-234 in orange, D2235-660,701-746 in purple, D3661-700,747-914 in cyan, D4915-1,074 in yellow, D51,075-1,273 in pink, and D61,274-1,424 in grey. **f-h**, SlaA glycans modelled with GlycoSHIELD in the assembled S-layer. (f) shows the extracellular view; (g) shows the intracellular view; (h) shows insets of (f) at higher magnification without (left) and with (right) glycans. Glycans fill gaps unoccupied by the protein and significantly protrude in the lumen of the triangular and hexagonal pores. Scale bars in (a-d and f-h), 10 nm; in (e), 20 Å.

Six SlaA monomers assemble around a hexagonal pore with 48 Å in diameter (glycans not included) (Fig. 4a). The D1 domains of these six monomers project into and define the shape of the hexagonal pore, together with D3 and D4 domains. The triangular pores that surround the hexagonal pores have a diameter of ∼85 Å and are defined by the D2, D4, D5 and D6 domains of three SlaA molecules (Fig. 4e). The D3 domain of each monomer overlaps with the D4 domain of the following monomer along the hexagonal ring in a clockwise fashion. D5 and D6 domains of each SlaA subunit project towards the cytoplasmic membrane. Two SlaA monomers dimerise through the D6 domains, with each SlaA dimer spanning two adjacent hexagonal pores (Fig. 4 b,e). Thus, protein-protein interactions between two adjacent hexagonal pores occur through the dimerizing D6 domains of each SlaA dimer and the D2 domains of overlapping SlaA monomers. The SlaA dimer includes an angle of 160° between the two monomers, and a total length of 420 Å (Supplementary Fig. 14a). While the SlaA was not resolved as a dimer in our single particle analysis, we could confirm these dimers in tomograms of negatively stained S-layers (Supplementary Fig. 15), which show similar dimensions and structure as in our assembly model.

**Figure 5.**
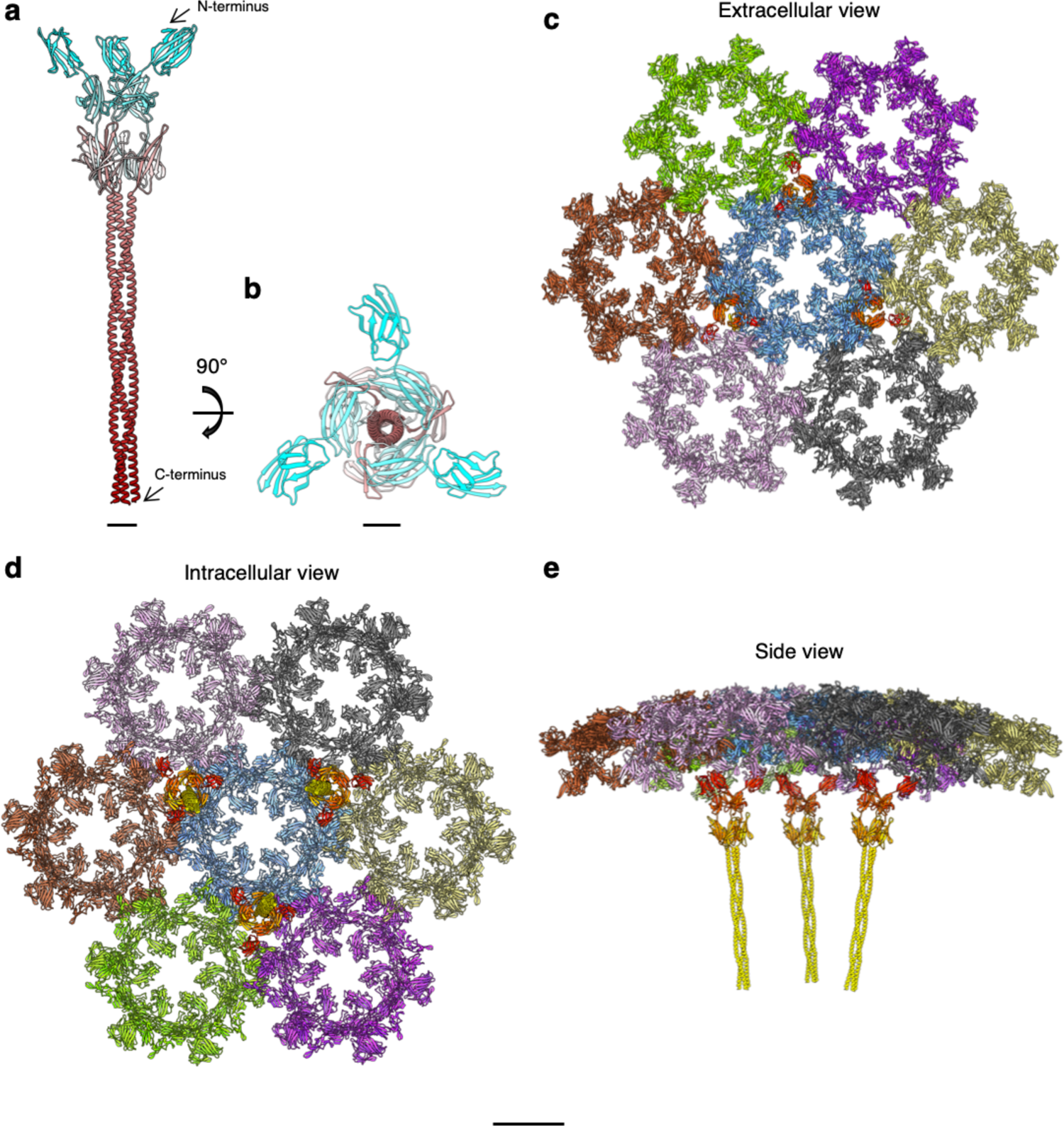
*S. acidocaldarius* S-layer assembly. **a** and **b**, SlaB trimer (ribbon representation, N terminus, cyan; C-terminus, maroon) as predicted by Alphafold v2.2.0. **c-d**, ribbon representation of the assembled SlaA and SlaB components of the S-layer. (c), (d), and (e) are extracellular, intracellular and side views, respectively. SlaA proteins around each hexagonal pore are shown in different colours. SlaB trimers are shown in orange tones (N-termini are in darker shades and C-termini in lighter shades). Scale bar, (a) and (b), 20 Å; (c-e) 10 nm.

Modelling of glycan shields in the assembled structure showed that glycans fill large gaps seen between SlaA’s globular domains and significantly protrude into the lumen of the triangular and hexagonal pores (Fig. 4 f-h). Most accessible sites in isolated SlaA monomers are still accessible in the assembled structure (Supplementary Fig. 8). Reduction of glycan conformational freedom is overall small between isolated and assembled SlaA monomers. Interestingly, the glycoshields appear to delineate protein-protein interfaces, which may guide the self-assembly of the S-layer. A glycan-guided assembly mechanism has been suggested for the assembly of cadherins in the desmosome^39^.

The cryoET map of the exosome S-layer contained limited information on SlaB. This is likely due to the previously reported flexibility of this protein^23^, meaning that it was mostly averaged out during STA. To get a handle on the structure of the entire S-layer, we used Alphafold v2.2.0^29^ and SymmDock^40^ and predicted the monomeric and trimeric SlaB structure. The resulting model is corroborated by the partially-resolved SlaB density visible in our STA map at low threshold values (Supplementary Fig. 12 g-i).

The SlaB structure is predicted to consist of three N-terminal β-sandwich domains and a 132 amino acid long C-terminal α-helix (Fig. 5a,b; Supplementary Fig. 16). The domain architecture and trimeric arrangement of SlaB agree with the sequence-based molecular modelling described by Veith *et al*.^21^ The TMHMM – 2.0 server predicted the N-terminal amino acids 448-470 as transmembrane helix. The hydrophobicity plot confirms a hydrophobic region corresponding to the predicted trans-membrane helix (Supplementary Fig. 17 a,e). The protein is predicted to have 14 N-glycosylation sites, of which six are located along the C-terminal α-helix (Supplementary Fig. 17b-d). The electrostatic surface potential calculated at pH 4 shows that the C-terminal α-helix is mostly neutral (Supplementary Fig. 17 f). In contrast, the three β-sandwich domains have greater electrostatic potential. While D2 is mostly positive, D3 carries distinct negatively charged patches (Supplementary Fig. 17f). These patches may be crucial in electrostatic interactions between SlaB’s D3 domain and the mainly positively charged SlaA.

By combining SPA and STA with structural predictions, we were able to build a complete *S. acidocaldarius* S-layer model (Fig. 5 c-e). In the assembled lattice, SlaB trimers occupy alternating triangular pores around each hexagonal pore^23^. The SlaB trimer has a tripod-like shape, with its long axis perpendicular to the membrane plane and that formed by SlaA. Three Ig-like domains branch away from the trimer’s symmetry axis and face the SlaA canopy, whereas three α-helices form a coiled coil and are partially inserted into the cytoplasmic membrane. The lattice is a ∼35 nm thick macromolecular assembly, in which each SlaB trimer interacts with three SlaA dimers. This interaction is mediated by the positively charged D6 dimerizing domains of SlaA and the negatively charged N-terminal Ig-like D3 domains of SlaB.

## Discussion

The *Sulfolobales* S-layer lattice stands out from other S-layers because it is a two-component lattice. The most likely advantage of such system is the ability of the cell to modulate the outermost component of the S-layer (SlaA) in response to environmental challenges, while conserving the membrane anchor (SlaB). In 2019, we reported on the structure of the *S. acidocaldarius* S-layer obtained from STA on whole cells and isolated S-layer sheets^23^. With the new information provided in the current study, we are able to improve on the model we proposed previously. The new data confirm the overall p3 S-layer lattice topology, in which the unit cell is constituted by one SlaB trimer and three SlaA dimers (SlaB_3_/3SlaA_2_). Each SlaB trimer occupies alternating triangular pores and each SlaA dimer spans two adjacent hexagonal pores. In agreement with our previously published model, we observe SlaB in alternating triangular pores. Because each SlaB monomer interacts with the dimerization domains of SlaA dimers, the SlaB trimer occupancy of all triangular pores would likely be unfavourable due to steric hindrance. Additionally, alternating SlaB throughout the array would reduce the protein synthesis costs for this protein by 50%. SlaB trimers occupying every second triangular pore also effectively create an S-layer with a variety of pore sizes, modulating the exchange of molecules with the environment.

Using exosomes and a new image processing approach, we were able to improve the resolution and eliminate the missing wedge in our subtomogram average of the *S. acidocaldarius* S-layer. The new map enabled us to build a revised model of the *S. acidocaldarius* S-layer assembly (Fig. 4 and 5). Here, the SlaA dimer (Supplementary Figure 14 a) spans an angle of 160° and extends over 42 nm, instead of 23 nm, as previously proposed^23^. The increased length is largely a result of the unexpected positioning of domains D5 and D6, which were previously not accounted for (Supplementary Fig. 14).

SLPs of extremophilic archaea generally show a high degree of glycosylation, potentially aiding their survival in extreme environments^41^. SlaA is predicted to contain 31 N-glycosylation sites^30^ and the SlaA_30-1,069_ cryoEM map showed 19 clear densities corresponding to N-glycosylation sequons. The cryoEM map contained densities for the complete hexasaccharide^30, 31^ on the SlaA surface, as well as various glycan intermediates. We cannot rule out the possibility that our cryoEM map could not resolve the complete hexasaccharide on all sequons due to the flexibility of the glycans. In any case, the presence of a heterogeneous family of glycans has also been reported by Peyfoon *et al*.^30^, who used nano-LC-ES-MS/MS to analyse the structure of the glycans linked to the C-terminus portion of SlaA (residues 961-1,395) and described a heterogenous degree of glycosylation including all intermediates from monosaccharide to complete hexasaccharide. The presence of a heterogeneous family of glycans has also been shown, for example, in the SLP of *H. volcanii*^42^ and the archaellum of *Methanothermococcus thermolithotrophicus*^43^. In archaea the final step in protein glycosylation is catalysed by the oligosaccharyl transferase AglB^44^. The enzyme is promiscuous, meaning that AglB is able to load glycans of variable length on the lipid carrier^45^. While AglB is essential for the viability of *S. acidocaldarius*^44^, it remains to be determined whether the heterogenous composition of its glycans is to be attributed to AglB loading glycan precursors onto SlaA and/or glycan hydrolysis due to the harsh environmental conditions.

Metal ions are often bound to SLPs and have recently been demonstrated to play a crucial role in S-layer assembly and cell-surface binding^15, 16, 46–50^. In the bacterium *C. crescentus*, whose S-layer has been investigated in detail, Ca^2+^ are essential for intra- and inter-molecular stability of the S-layer lattice^15, 49^. Moreover, analogous results have been obtained for the S-layer of *Geobacillus stearothermophilus*^16^. The SLP of the archaeon *H. volcanii* has also been recently confirmed to bind cations^14^. The *S. acidocaldarius* S-layer is no exception and its assembly is a Ca^2+^-dependent process^23^. Interestingly, the SlaA_30-1,069_ cryoEM map did not reveal any anomalous densities that could be attributed to ions. It is therefore possible that cations are harboured in the D5 and D6 domains that were not resolved, and / or at the protein-protein interfaces within the assembled lattice.

In a recent work, von Kügelgen *et al*. have presented the structure of the *H. volcanii* S-layer^14^. Therefore, the *H. volcanii* and *S. acidocaldarius* S-layers are currently the only two archaeal S-layers for which complete atomic models are available. *H. volcanii* is a halophilic archaeon of the Euryarchaeota phylum. As the *S. acidocaldarius* S-layer, the *H. volcanii* lattice also exhibits a hexagonal symmetry, but different architecture. The *H. volcanii* S-layer is constituted by a single glycosylated SLP named csg. SlaA (1,424 residues) and Csg (827 residues) both consist of six domains (Supplementary Fig. 18 b). However, while all csg domains adopt Ig-like folds, SlaA is built up from domains of more complex topology. In csg, the domains are arranged linearly, whereas SlaA adopts an extended Y-shape (Supplementary Fig. 18 a,b). Ig-like domains are widespread among SLPs in different archaeal phyla, including the order Sulfolobales^14^. In fact, the SlaA protein of *Metallosphaera sedula* is predicted to consist of seven Ig-like domains (Supplementary Fig. 18 d)^14^. The different domain architecture that we observe for *S. acidocaldarius* SlaA highlights the great divergence of S-layers among microorganisms.

Assembled csg forms hexagonal (13 Å)/pentameric (6 Å) and trimeric (10 Å) pores much smaller than the hexagonal (48 Å) and trimeric (85 Å) pores of the *S. acidocaldarius* lattice. In both cases, the pore size is further reduced by glycans projecting into the pores. The glycans could regulate the permeability of the S-layer in a fashion similar to the hydrogel regulating the permeability of the nuclear pore complexes^51^. It is currently unknown, which evolutionary parameters resulted in species specific S-layer pore sizes. It may be speculated that, for example, these pores have co-evolved with and adapted their size according to certain secreted protein filaments, such as pili. *Sulfolobus* produces four such filaments - archaella^52^, A-pili^53^, UV-inducible pili and threads^54^. Of these four filaments, only threads, with a diameter of ∼40 Å, would be able to pass through the hexagonal pores of the S-layer without the need for partial S-layer disassembly. It is thus tantalising to speculate that the hexagonal S-layer pores have evolved to accommodate threads, perhaps as a scaffold for their assembly.

S-layers are intrinsically flexible structures as to encapsulate the cell entirely. In the case of *H. volcanii*, csg assembles around hexameric as well as pentameric pores on the surface of both exosomes and whole cells^14^. Such pentameric defects confer enough flexibility to the array to encase the cell in areas of low and high membrane curvature. Interestingly, we did not observe an analogous phenomenon for the *S. acidocaldarius* S-layer on whole cells or exosomes. However, symmetry breaks have been observed on S-layers isolated from whole cells at the edges where the lattice changes orientation^55^. Furthermore, additional flexibility may be provided by the SlaA dimeric interface, as well as by loop regions linking the SlaA domains. In fact, only single loops link D1-D2, D3-D4, D4-D5 and D5-D6. While the reciprocal position of D3-D4 is stabilised by the disulphide bond (Cys_677_-Cys_1,017_), the loops connecting D1-D2, D4-D5 and D5-D6 may allow the flexibility necessary for SlaA to be incorporated in this highly interwoven, yet deformable protein network.

Electrostatic interactions are critical for proper protein folding and function. Moreover, changes in surface charge have been shown to affect protein-protein interactions. Particularly, the pH plays a key role in determining the surface charge of proteins due to polar amino acid residues on the protein surface^35, 36^. Remarkably, SlaA_30-1,069_ proved stable over a vast pH range and its tertiary structure remains virtually unchanged. Furthermore, the surface net charge of SlaA shifts from positive to negative from pH 4 to pH 10. In view of this, we hypothesise that such transition in surface charge and not the protein unfolding is responsible for SlaA disassembly at pH 10. Considerations regarding the pH stability of SlaA_30-1,069_ can be extended to the entirety of the protein using pH stability predictions, which suggest virtually no difference in pH-dependent protein stability across ionic strength and pH values for both SlaA_30-1,069_ and SlaA (Supplementary Fig. 19a-d). This suggests that domains D5 and D6 equally do not unfold at alkaline pH. Analogous predictions of protein stability were obtained for SlaB (Supplementary Fig. 19e,f), where the net charge is slightly positive across pH 2-8. For comparison, we ran the same predictions on the *C. crescentus* and *H. volcanii* S-layer proteins RsaA and csg, respectively (Supplementary Fig. 20). Among SlaA, SlaB, RsaA and csg, we observe that SlaA and SlaB are expected to be the most stable at different pH values. Notably, csg is most stable at acidic pH and progressively less so at pH neutral and alkaline. This prediction is confirmed by experimental data^56^, which additionally showed pH-dependent protein folding rearrangements and protein unfolding. It is to be considered that this prediction does not <u>include</u> glycosylation^57^, which enhances S-layer stability, especially in the case of Sulfolobales^41, 44, 58, 59^. The resilience of SlaA at temperature and pH shifts can likely be attributed to two main factors: the high glycosylation level, and the fact that ∼ 56% of SlaA_30-1,069_ has a defined secondary structure, which allows the formation of intramolecular bonds^60^.

S-layers are often necessary for the survival of microorganisms in nature but can also be of great interest for synthetic biology. Therefore, a greater understanding of their structural details will strongly aid their nanotechnological uses, which have already shown remarkable potential in biomedical and environmental applications^61–64^.

## Methods

### *S. acidocaldarius* strains and growth conditions

Cells of *S. acidocaldarius* strain MW001 were grown in basal Brock medium* at pH 3^65^ as previously described^23^. Briefly, cells were grown at 75 °C, 150 rpm, until an OD600 of >0.6 was reached. Cells were then centrifuged at 5,000 g (Sorval ST 8R) for 30 min at 4 °C. The cell fraction was stored at −20 °C for S-layer isolation, whereas the supernatant was stored at 4 °C for exosomes isolation. *Brock media contain (per l): 1.3 g (NH_4_)2SO_4_, 0.28 g KH_2_PO_4_, 0.25 g MgSO_4_ • 7H_2_O, 0.07 g CaCl_2_ • 2H_2_O, 0.02 g FeCl_2_ • 4H_2_O, 1.8 mg MnCl_2_ • 4H_2_O, 4.5 mg Na_2_B_4_O_7_ • 10H_2_O, 0.22 mg ZnSO_4_ • 7H_2_O, 0.05 mg CuCl_2_ × 2H_2_O, 0.03 mg NaMoO_4_ • 2H_2_O, 0.03 mg VOSO_4_ × 2H_2_O, 0.01 mg CoSO_4_ • 7H_2_O, and 0.01 mg uracil.

### S-layer isolation and disassembly

The S-layer isolation and disassembly were performed as previously described^66^. Briefly, frozen cell pellets from a 50 ml culture were incubated at 40 rpm (Stuart SB3) for 45 min at 37 °C in 40 ml of buffer A (10 mM NaCl, 1 mM phenylmethylsulfonyl fluoride, 0.5% sodium lauroylsarcosine), with 10 μg/ml DNase I. The samples were pelleted by centrifugation at 18,000 × g (Sorvall Legend XTR) for 30 min and resuspended in 1.5 ml of buffer A, before further incubation at 37 °C, for 30 min. After centrifugation at 14,000 rpm for 30 min (Sorvall ST 8R), the pellet was purified by resuspension and incubation in 1.5 ml of buffer B (10 mM NaCl, 0.5 mM MgSO_4_, 0.5% SDS) and incubated for 15 min at 37 °C. To remove SlaB from the assembled S-layers, washing with buffer B was repeated <u>three more times</u>. Purified Sla-only S-layers were washed once with distilled water and stored at 4 °C. The removal of SlaB was confirmed by SDS/PAGE analysis. S-layers were disassembled by increasing the pH to 10 with the addition of 20 mM NaCO_3_ and 10 mM CaCl_2_ and incubated for 2 hours at 60 °C, 600 rpm (Thermomixer F1.5, Eppendorf).

### SlaA purification

After disassembly the sample containing SlaA was further purified using gel filtration chromatography. A total of 100 μl containing 10 mg/ml of disassembled protein were loaded onto a Superdex 75 Increase 10/300 GL (GE Healthcare) using 300 mM NaCl for elution. At the end of the run, the fractions containing SlaA were dialysed against 30 mM acetate buffer (0.1 M CH₃COOH, 0.1 M CH_3_COONa) at pH 4, 150 mM Tris-HCl at pH 7, or 20 mM NaCO_3_ at pH 10, with the aim to compare the SlaA protein structure at different pH values. The purity of the fractions was assessed by SDS/PAGE analysis and negative staining with 1% uranyl acetate on 300 mesh Quantifoil copper grids with continuous carbon film (EM Resolutions).

### CryoEM workflow for single particle analysis (SPA)

#### Grid preparation

The purified SlaA samples at pH 4 and 10 (3 μl of ∼0.1 mg/ml) were applied on 300 mesh copper grids with graphene oxide-coated lacey carbon (EM Resolutions) without glow discharge. Grids were frozen in liquid ethane using a Mark IV Vitrobot (Thermo Fisher Scientific, 4 °C, 100 % relative humidity, blot force 6, blot time 1 sec) with Whatman 597 filter paper. The purified SlaA at pH 7 was applied on glow discharged R 1.2/1.3 300 mesh copper grids with holey carbon. The freezing procedure was kept the same as for the samples at pH 4 and 10 besides the blot time 2 sec.

#### Data collection

Micrographs were collected on a 200 kV FEI Talos Arctica TEM, equipped with a Gatan K2 Summit direct detector using the EPU software (Supplementary Table 1). Data were collected in super-resolution at a nominal magnification of 130,000x with a virtual pixel size of 0.525 Å at a total dose of ∼60 e^-^/Å^2^. A total of 3,687 movies (44 fractions each), 3,163 movies (44 fractions each), and 5,046 movies (60 fractions each), with a defocus range comprised between −0.8 and −2.4 μm, were collected for samples at pH 4, pH 7 and pH 10, respectively.

#### Image processing

Initial steps of motion correction (MotionCor 2^67^) and CTF estimation (CTF-find 4^68^) were performed in Relion 3.0^34^ and Relion 3.1^25^ for datasets at pH 4 and 7, whereas Warp^69^ was used for the pH 10 dataset. Further steps of 2D and 3D classification, refinement, CTF refinement and polishing were performed using Relion 3.1. For a detailed workflow of the three datasets see Supplementary Fig. 1, 9a,b and 10. The refined maps were post-processed in Relion 3.1 as well as using DeepEMhancer^70^. The produced maps had a resolution of 3.1 Å, 3.9 Å and 3.2 Å at pH 4, 7 and 10, respectively, by gold-standard FSC 0.143.

#### Model building and validation

The SlaA atomic model was built *de novo* using the cryoEM map at pH 4 in Buccaneer^71^, refined using REFMAC5^72^ and rebuilt in *Coot*^73^. The glycans were modelled in *Coot* with the refinement dictionary for the unusual sugar 6-sulfoquinovose prepared using JLigand^74^. This atomic model was then positioned into the cryoEM maps at pH 10 and pH 7 using ChimeraX^75^ and refined using REFMAC5 and *Coot*. All models were further refined using Isolde^76^ and validated using Molprobity^77^ in CCP4^78^.

### Exosome isolation

*S. acidocaldarius* exosomes were isolated from the supernatant obtained after cell growth. The procedure was adapted from Ellen *et al*., 2009^37^. The supernatant was split into 8 fractions and exosomes were pelleted in two runs of ultracentrifugation (Optima LE-80K, Beckman Coulter) at 125,000 g for 45 min at 4 °C. The pellet was resuspended in 2 ml (per fraction) of the supernatant and ultracentrifuged (Optima MAX-TL, Beckman Coulter) at 12,000 rpm (TLA55 rotor, Beckman Coulter) for 10 min at 4 °C. The pellet (containing intact cells and cell debris) was discarded, and the supernatant was ultracentrifuged (Optima MAX-TL, Beckman Coulter) at 42,000 rpm (TLA55 rotor, Beckman Coulter) for 90 min at 4 °C. The pellet containing the isolated exosomes was resuspended in MilliQ water at a concentration of 15 mg/ml. The purity of the sample was assessed by negative staining with 1% uranyl acetate on 300 mesh Quantifoil copper grids with continuous carbon film (EM Resolutions).

### CryoEM workflow for subtomogram averaging

#### Grid preparation

The isolated exosomes were mixed 1:1 with 10 nm colloidal gold conjugated protein A (BosterBio) and 3 μl droplets were applied four times on glow discharged 300 mesh Quantifoil copper R2/2 grids (EM Resolutions). The grids were blotted with 597 Whatman filter paper for 4 sec, using blot force 1, in 95 % relative humidity, at 21 °C, and plunge-frozen in liquid ethane using a Mark IV Vitrobot (FEI).

#### Data collection

Micrographs were collected on a 200 kV FEI Talos Arctica TEM, equipped with a Gatan K2 Summit direct detector using the Tomo 4 package. Tilt series were collected in super-resolution at a nominal magnification of 63,000 x with a virtual pixel size of 1.105 Å at a total dose of ∼83 e-/Å2. The tilts were collected from –20 deg to 60 deg, in 3 degree steps (2 fractions per tilt). A defocus range between −4 and −6 μm was used for the collection. A total of 28 positions were collected.

#### Electron cryotomography and subtomogram averaging

Tilt series motion correction was performed using the IMOD^79^ program alignframes. IMOD was also used for the tomogram reconstruction. Initial particle picking on all 28 tomograms was performed using seedSpikes and spikeInit as part of the PEET software package^80^ with a total of 12,010 particles picked. For initial subtomogram averaging, the picked particles were CTF corrected and extracted using the Relion STA pipeline^81^. 2D classification, initial model generation, 3D classification and initial refinements were all performed using Relion 3.1^25^. A resolution of 16.1 A was reached using 1,313 particles and C3 symmetry.

For high resolution averaging, emClarity^38^ was used. After 3D CTF correction and subregion selection, template matching was performed at a 4x binning, using a Relion-generated map centred on the hexagonal pore with a total of 3,210 particles picked. Initial averaging began with C1 symmetry at 3x binning, with iterative tilt series refinement followed by a reduction in binning to 2x and, finally, unbinned particles. During the unbinned processing, a symmetry of C3 was applied. The final map achieved a gold-standard resolution of 8.9 Å (FSC 0.143) using 3,182 particles.

### Structure analysis and presentation

The electrostatic potential of the protein was derived using APBS (Adaptive Poisson-Boltzmann Solver)^82^ based on the PARSE force field for the protein as available through PDB2PQR^83^. Where available, the charges of the glycans were assigned based on the GLYCAM force field^84^; charges of the hydrogens were combined with their central heavy atom. The charge assignment depends on the bonding topology, i.e. occupied linkage positions. Supplementary Table 2 summarizes the mapping of residue from the structure file to GLYCAM residue names. For residue SMA, charge assignments are not available from the GLYCAM force field; these were derived based on RESP calculations conducted for the methoxy-derivatives on the HF/6-31G*//HF/6-31G* level of theory and employing a hyperbolic restraint equal to 0.010 in the charge fitting step^85, 86^. The total charge of the newly derived residue was constrained to − 0.8060 e and −1 e for the 1-substituted and 1,4-substituted SMA (referred to as SG0 and SG4 in Supplementary Table 3 & 4), respectively, in agreement with the conventions of the GLYCAM force field. In assembling the final charge assignment, the charge of the linking ND2 atom of the glycosylated Asn residues of the protein were altered to compensate for the polarization charge of the attached saccharide unit. The electrostatic charge was visualised using VMD^87^ (http://www.ks.uiuc.edu/Research/vmd/).

The structure of *S. acidocaldarius* SlaA was visualised using UCSF Chimera^88^, Chimera X v.1.3 and v1.4^75^, and Pymol. The structural domains of SlaA were assigned using SWORD^26^.

Heatmaps for net charge, and pH and ionic strength-dependent protein stability were obtained using Protein-Sol (https://protein-sol.manchester.ac.uk/)^57^. For SlaB (Supplementary Fig. 20 and 21) the signal-peptide was predicted using InterPro^89^, the trans-membrane region was predicted using TMHMM – 2.0^90^, the N-glycosylation sites (sequons N-X-S/T) were predicted using GlycoPP v1.0^91^.

### Molecular dynamics simulations (MDS)

Conformation arrays of glycans were grafted on protein structure using GlycoSHIELD^33^. In brief, glycan systems (GlcNAc[2],Man[2],QuiS[1],Glc[1] N-linked to neutralised glyc-Asp-gly tripeptides) were modelled in CHARMM-GUI^92^ and solvated using TIP3P water models in the presence of 150mM NaCl and configured for simulations with CHARMM36m force fields^93, 94^. MDS were performed with GROMACS 2020.2 and 2020.4-cuda^95^ in mixed GPU/CPU environments. Potential energy was first minimized (steepest descent algorithm, 5,000 steps) and were equilibrated in NVT ensemble (with 1 fs time-steps using Nose-Hoover thermostat). Atom positions and dihedral angles were restrained during the equilibration, with initial force constants of 400, 40 and 4 kJ/mol/nm^2^ for restraints on backbone positions, side chain positions and dihedral angles, respectively. The force constants were gradually reduced to 0. Systems were additionally equilibrated in NPT ensemble (Parrinello-Rahman pressure coupling with the time constant of 5 ps and compressibility of 4.5 10^-5^ bar^-1^) over the course of 10 ns with a time step of 2 fs. Hydrogen bonds were restrained using LINCS algorithm. During the production runs, velocity-rescale thermostat was used and temperature was kept at 351K. Production runs were performed for a total duration of 3μs and snapshots of atom positions stored at 100 ps intervals.

Glycan conformers were grafted using GlycoSHIELD with a distance of 3.25 Å between protein α-carbons and glycan ring-oxygens. Glycan conformers were shuffled and subsampled for representation of plausible conformations on displayed renders. Graphics were generated with ChiemeraX^75^.

### Data availability

The SlaA atomic coordinates were deposited in the Protein Data Bank (https://www.rcsb.org/) with accession numbers 7ZCX, 8AN3, and 8AN3 for pH 4, 7 and 10, respectively. The electron density maps were deposited in the EM DataResource (https://www.emdataresource.org/) with accession numbers EMD-14635, EMD-15531 and EMD-15531 for pH 4, 7 and 10, respectively. Other structural data used in this study are: *H. volcanii* csg (PDB ID: 7PTR, http://dx.doi.org/10.2210/pdb7ptr/pdb), and *C. crescentus* RsaA ((N-terminus PDB ID: 6T72, http://dx.doi.org/10.2210/pdb6t72/pdb, C-terminus PDB ID: 5N8P, http://dx.doi.org/10.2210/pdb5n8p/pdb).

## Acknowledgements

We acknowledge Ufuk Borucu for help with data collection, and the GW4 Facility for High-Resolution Electron Cryo-Microscopy, funded by the Wellcome Trust (202904/Z/16/Z and 206181/Z/17/Z) and BBSRC (BB/R000484/1). We also acknowledge Alexander Neuhaus for assistance with data analysis. We thank IDRIS for the allocation of high-performance computing resources (allocations #2020-AP010711998 and #2021-A0100712343 to CH). For this project, L.G., B.D., M.M, MG and KS were funded by the European Research Council (ERC) under the European Union’s Horizon 2020 research and innovation programme (grant agreement No 803894). RC was funded by the University of Exeter and a Wellcome Trust Seed Award in Science (210363/Z/18/Z) awarded to VG as well as a Wellcome Trust Seed Award in Science (212439/Z/18/Z) awarded to BD. MS is supported by a Fonds zur Förderung der wissenschaftlichen Forschung Schrödinger fellowship (J4332-B28). Work in the laboratory of CH is supported by the Agence Nationale de la Recherche (grants #ANR-16-CE16-0009-01 and #ANR-21-CE16-0021-01). DK was funded by the Leverhulme Trust (RPG-2020-261).

## Author contributions

LG: Performed the research, provided methodology, wrote the manuscript.

MM: Provided methodology and performed research.

RC: Provided methodology.

KS: Provided methodology.

MG: Performed research.

LC: Performed research.

VG: Provided resources

DK: Provided methodology.

MS: Provided methodology.

CH: Provided methodology and performed research

MI: Performed research, provided methodology, wrote the manuscript.

BD: Provided funding, performed research, provided resources, conceptualised the research and wrote the manuscript.

## Supplementary Figures

**Supplementary figure 1.**
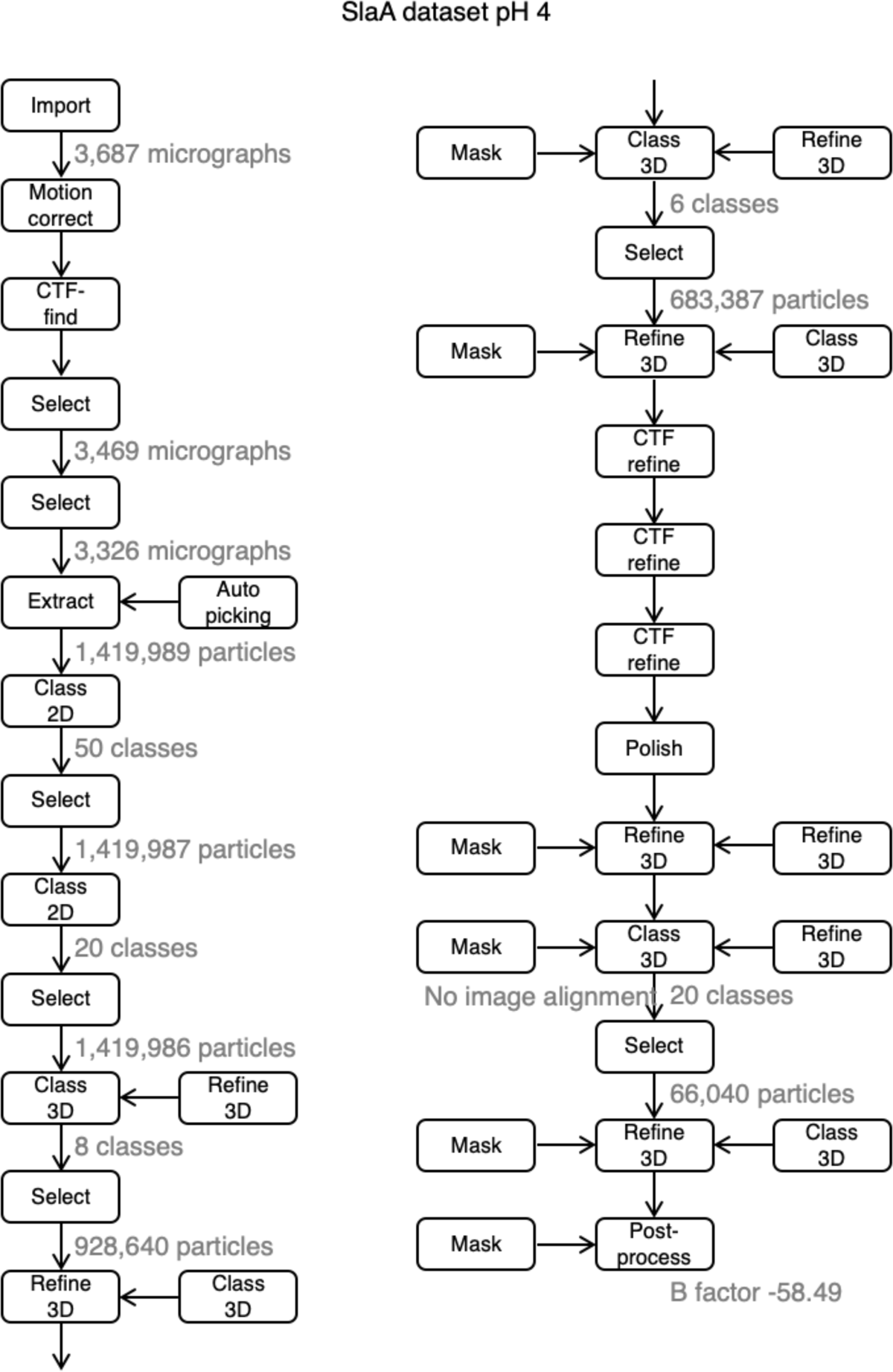
Relion processing workflow for pH 4 dataset.

**Supplementary figure 2.**
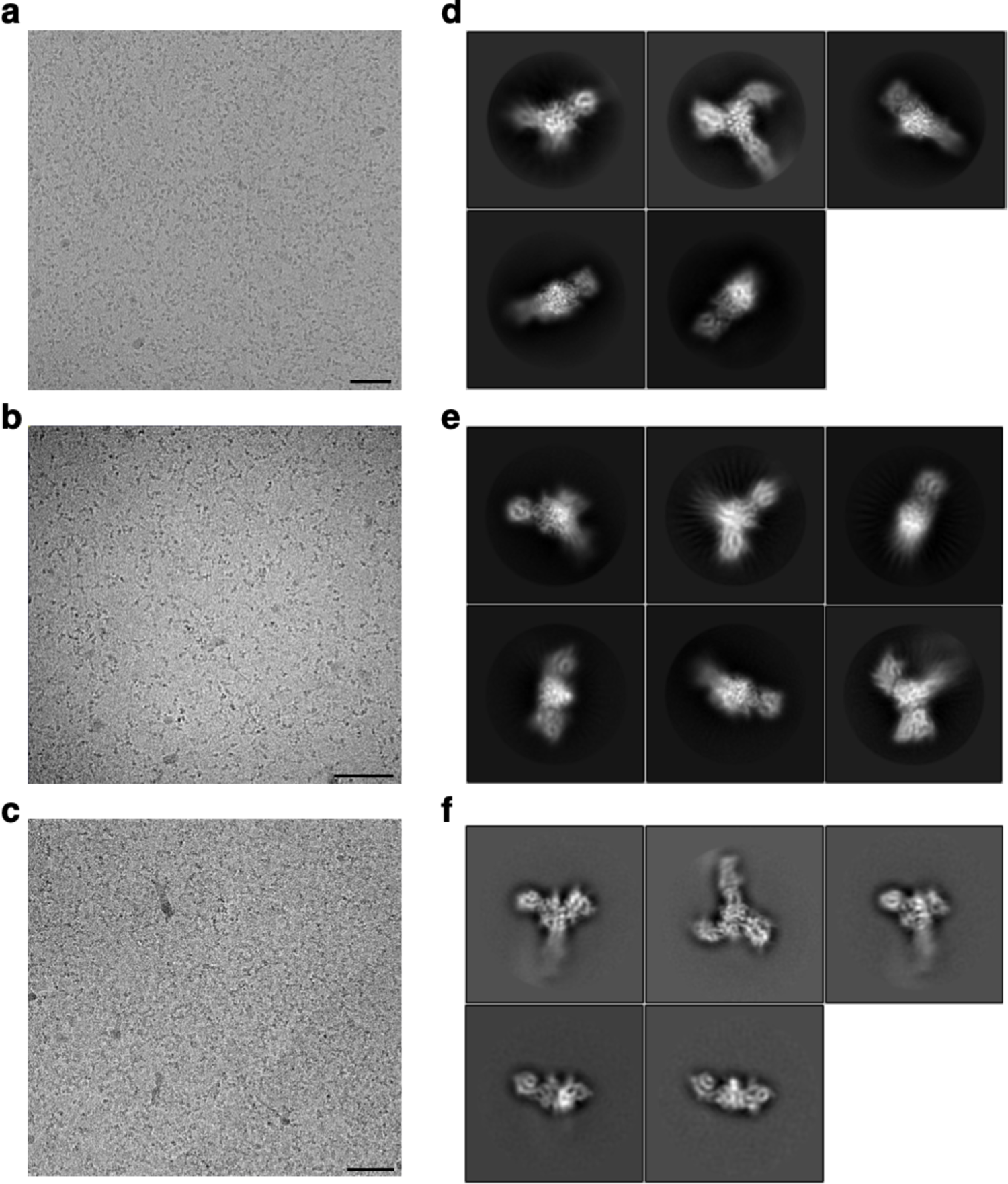
SlaA representative data. **a-c**, representative cryoEM micrographs (from a total of 3,687 for (a), 3,163 for (b) and 5,046 for (c)). **d-f**, 2D classification examples of *S. acidocaldarius* SlaA polished particles in Relion at pH 4 (a, d), pH 7 (b, e) and pH 10 (c, f). Scale bar; 50 nm.

**Supplementary figure 3.**
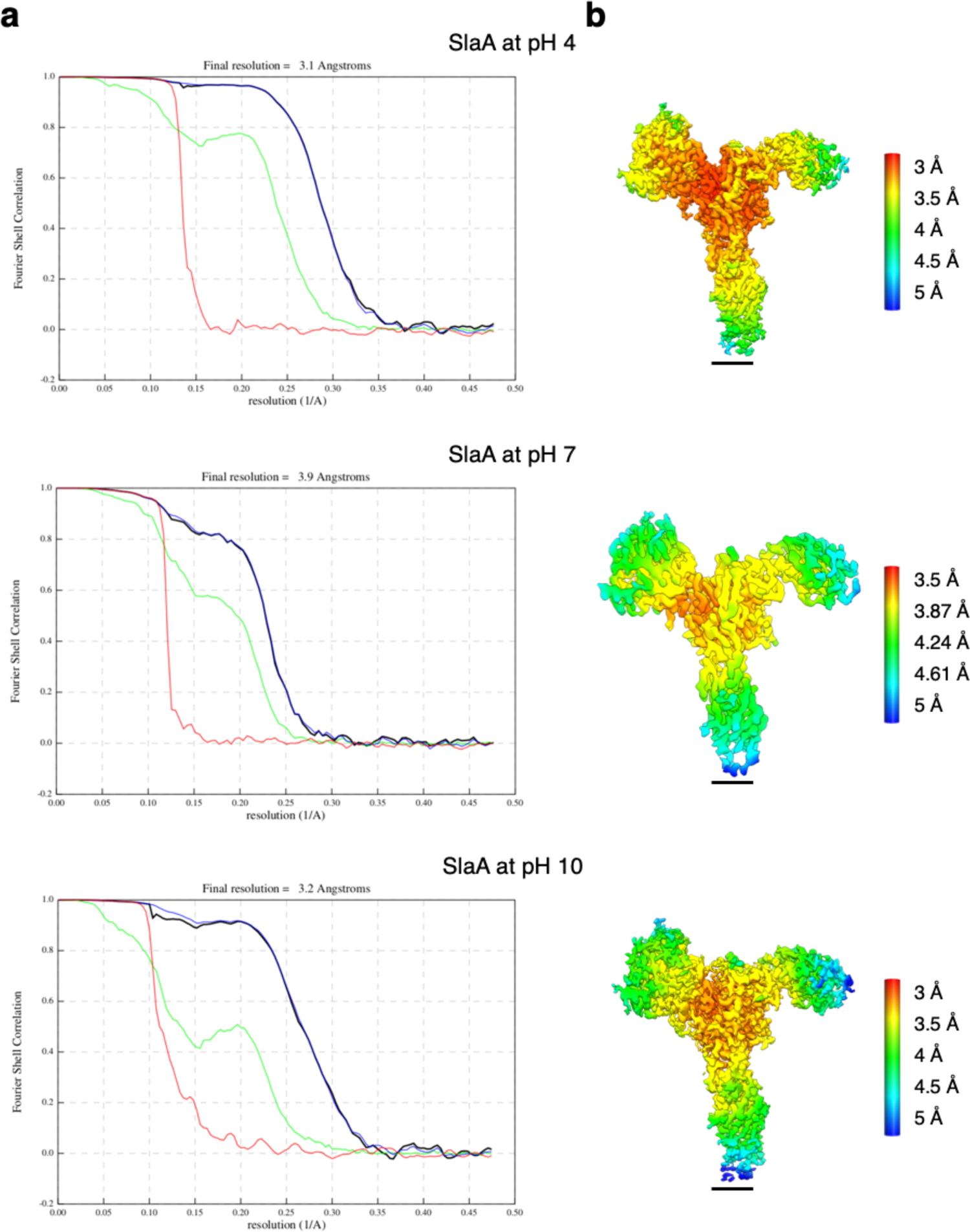
SlaA data quality in Relion. **a**, gold standard FSC and **b**, local resolution estimations for the SlaA map obtained at pH 4, 7 and 10. Red, phase randomised masked; green, unmasked; blue, masked; black, corrected. Scale bar, 20 Å.

**Supplementary figure 4.**
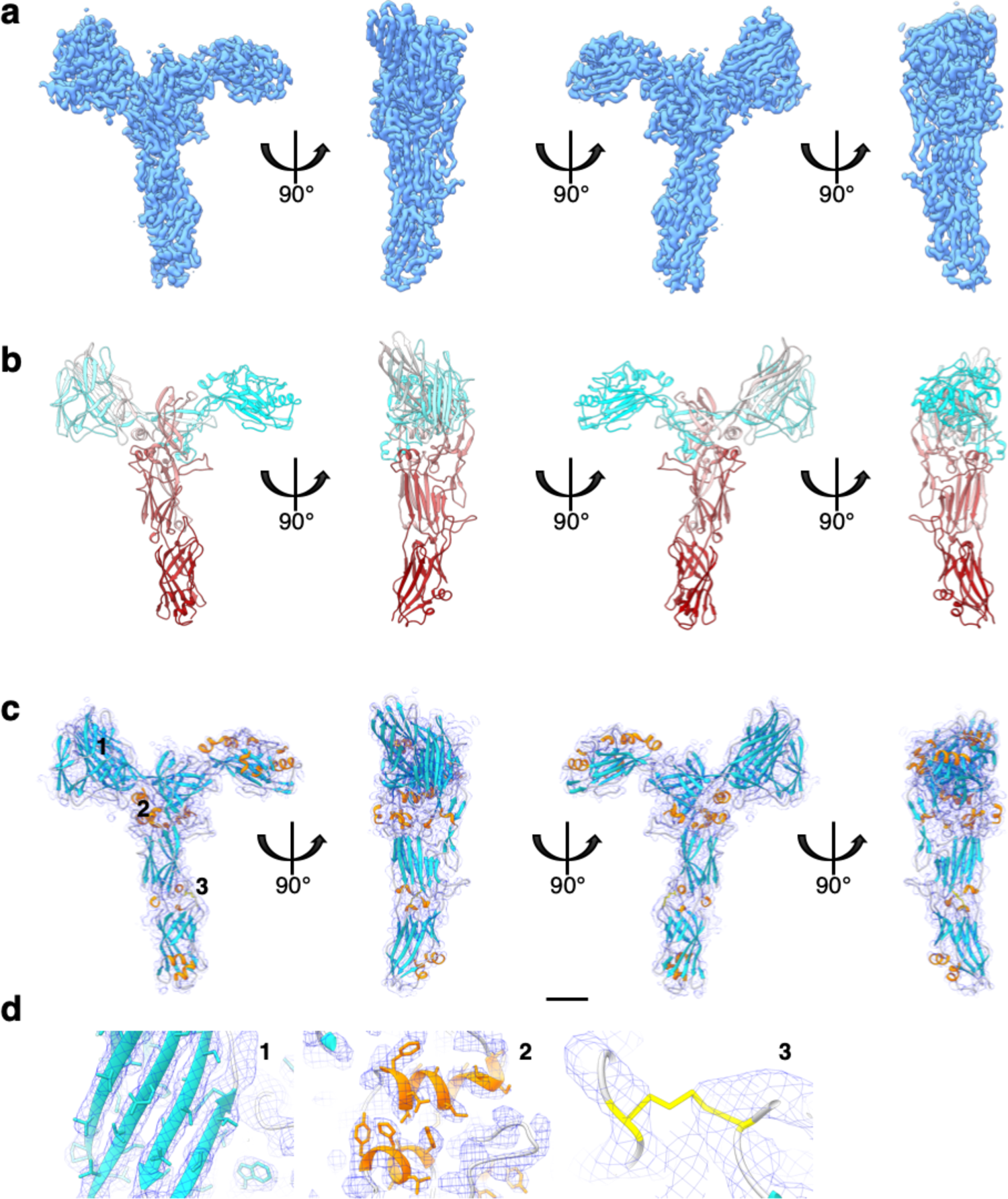
SlaA30-1069 cryoEM map and atomic model. **a**, SlaA30-1,069 cryoEM density map at 3.1 Å global resolution. **b**, atomic model of SlaA30-1,069 (ribbon representation, cyan-grey-maroon colours. N-terminus, cyan; C-terminus, maroon). **c**, fitting of the atomic model (ribbon representation) into the cryoEM density map (blue mesh). The loop regions are in grey, α-helices in orange, β-sheets in turquoise, and the disulphide bridge in yellow. **d**, close-ups of three example regions of β-sheets, α-helices and the disulphide bridge. Locations of the close-ups are labelled in (c) as 1, 2 and 3. Scale bar in (a-c), 20 Å.

**Supplementary figure 5.**
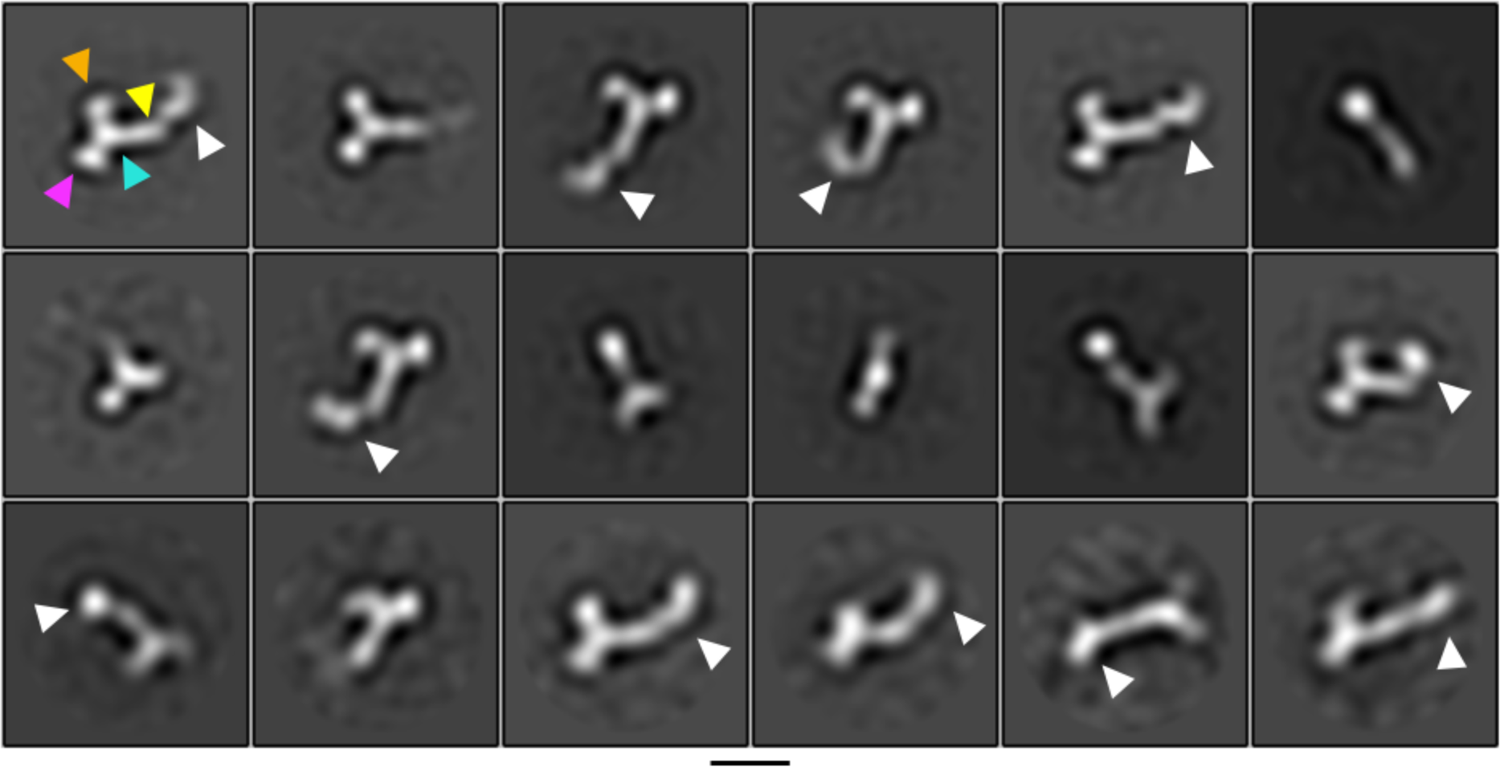
SlaA flexibility. 2D classification of negative staining micrographs of isolated SlaA. The white arrowheads point at domains D5 and D6 in different orientations, highlighting the mobility of these domains. The arrowheads in the first class highlight D1 (orange), D2 (purple), D3 (cyan), and D4 (yellow). Scale bar, 100 Å.

**Supplementary figure 6.**
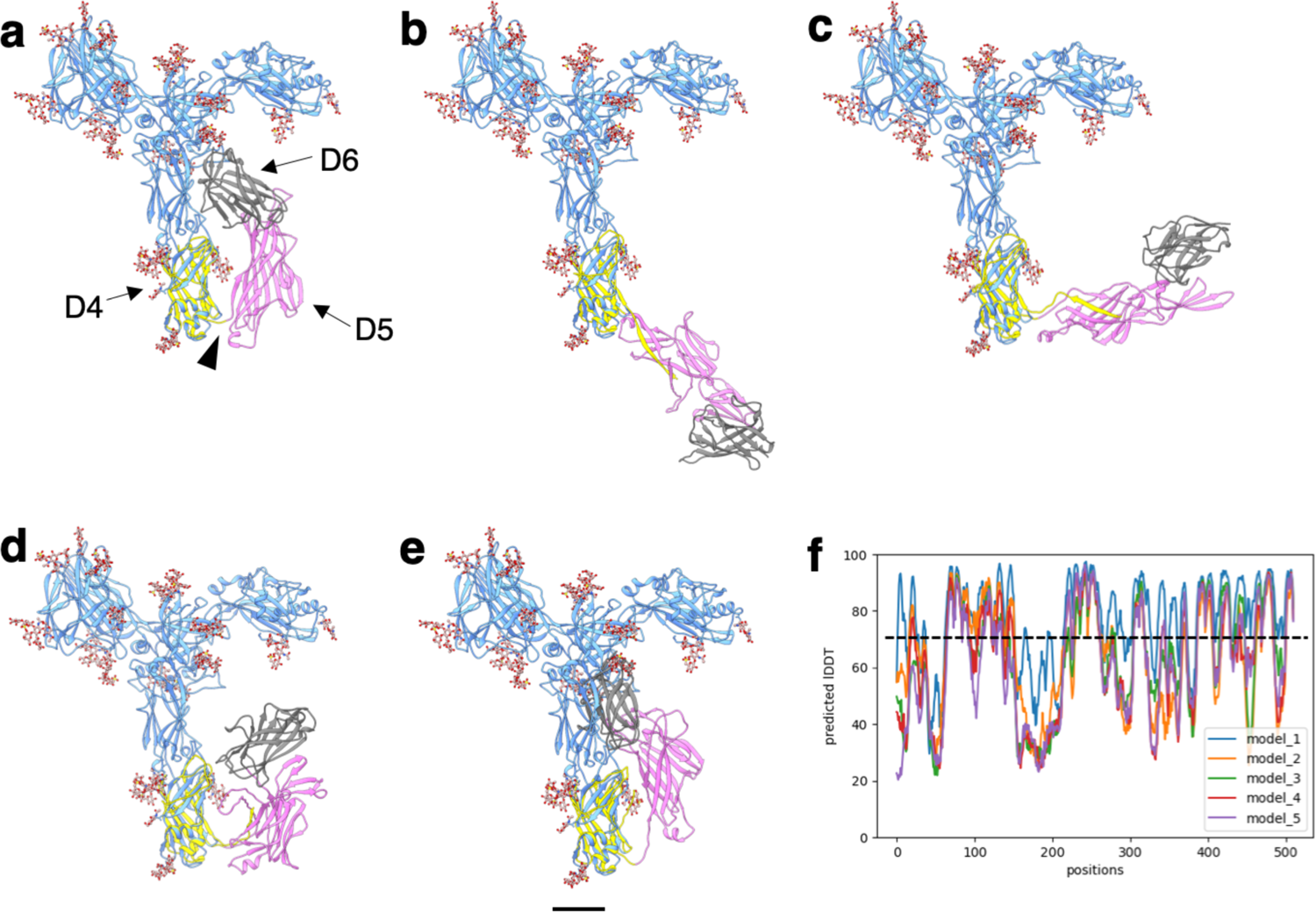
Five Alphafold predictions of SlaA914-1,424. **a-e**, SlaA30-1,069 is shown in ribbon representation and cornflower blue; glycans are in ball-stick representation and rosy brown. The Alphafold predictions are coloured according to domains, highlighting domains D5 (pink) and D6 (grey). Residues 914-1,069 (D4 in yellow) at the C-terminus of SlaA30-1,069 were included in the prediction to aid alignment between SlaA30-1,069 and D5-D6. The predicted D4 largely overlaps with the cryoEM structure of the SlaA30-1,069 N-terminus. The black arrowhead (a) indicates the intramolecular hinge loop. **f**, pLDDT (per-residue confidence score) plot showing the per-residue confidence metric of the predicted models. The dashed line marks the threshold of predicted LDDT=70, above which the structures are expected to be modelled with high confidence. Scale bar, 20 Å.

**Supplementary figure 7.**
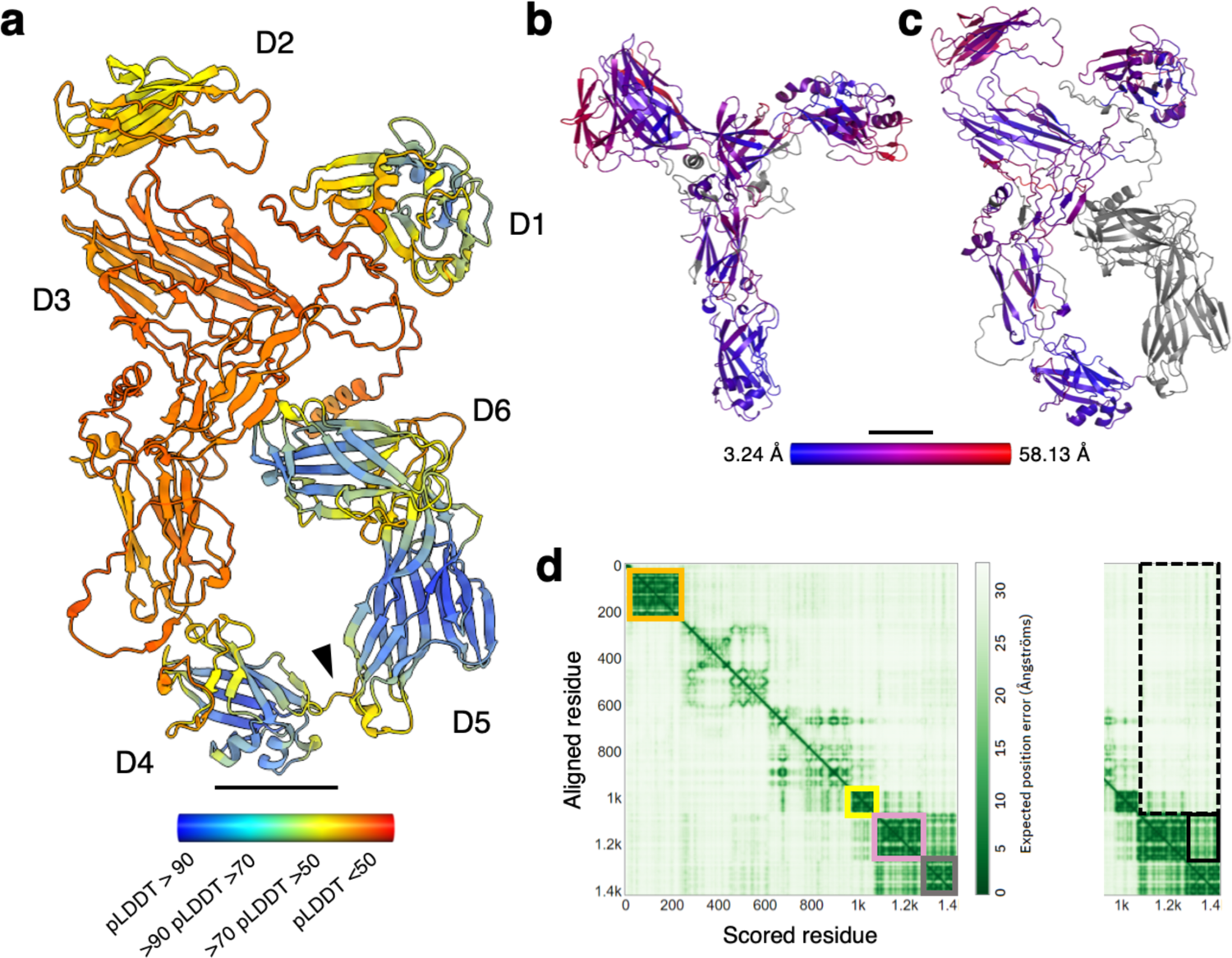
Alphafold v2.2.0 predictions of SlaA and comparison between SlaA30-1,069 solved by cryoEM and the Alphafold v2.2.0 prediction. a, SlaA as predicted by Alphafold v2.2.0. The ribbon is coloured by pLDDT (per-residue confidence score) where red indicates very low confidence and blue very high. Black arrowhead, intramolecular hinge loop. b and c, comparisons between SlaA30-1,069 solved by cryoEM (b) and the Alphafold v2.2.0 predictions (c). The ribbon is coloured by r.m.s.d. (root-mean-square deviation) showing smaller and larger deviations in blue and red respectively, with mean r.m.s.d. = 25.03 Å. Gray areas are not aligned. d, PAE (predicted aligned error) plot of the Alphafold v2.2.0 prediction. The boxed areas define predictions with high level of confidence: domains D1 (orange), D4 (yellow), D5 (pink) and D6 (grey). In the cut-out on the right the solid black line defines the confidence area for the reciprocal position of D5 and D6 (darker green, higher confidence), whereas the dashed line defines the confidence area for the reciprocal position of D5-D6 and the rest of the complex (lighter green, lower confidence). Scale bar, 20 Å.

**Supplementary figure 8.**
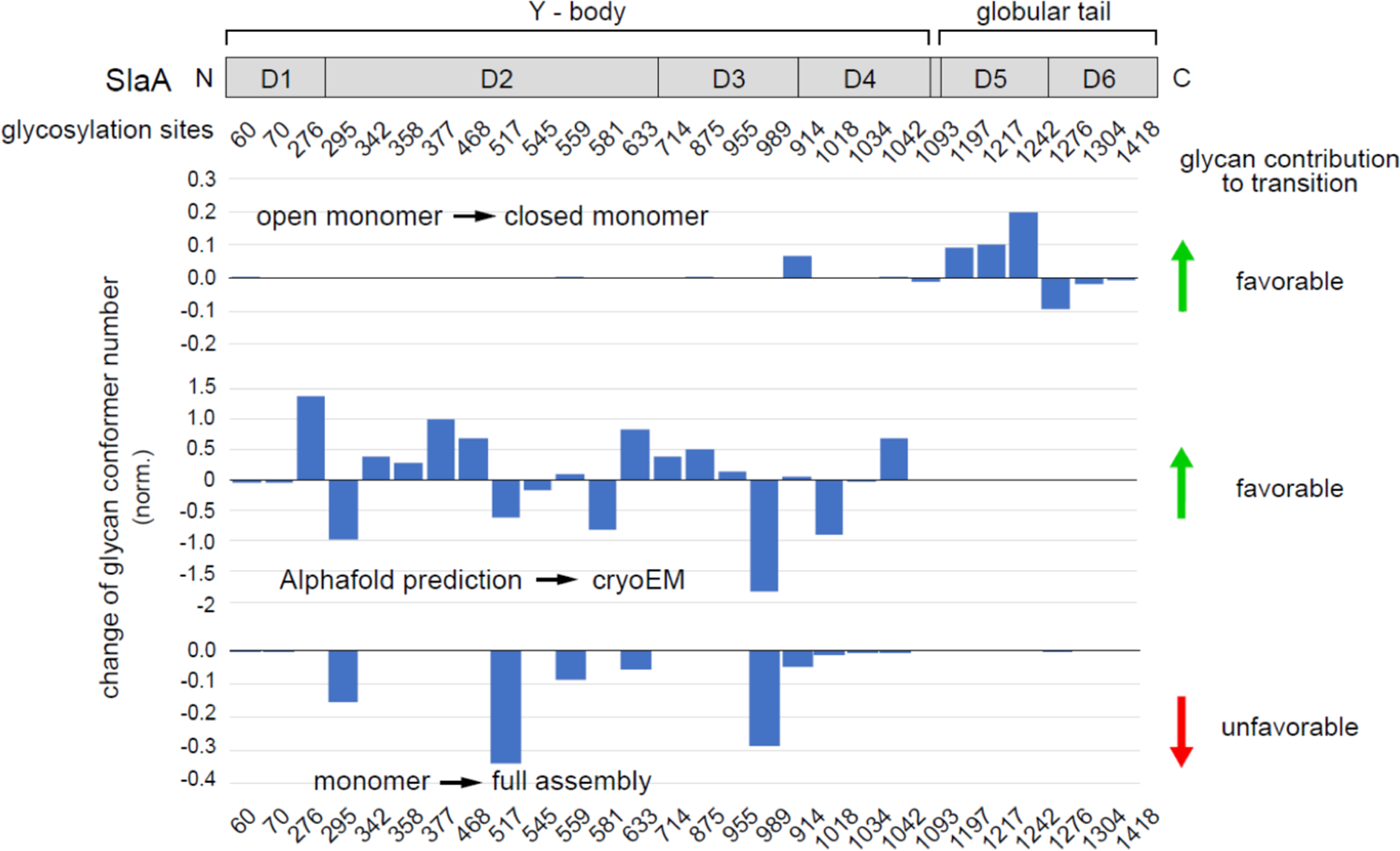
Entropic contribution of glycans to protein conformation. Position of N-glycosylated sites and globular domains on SlaA primary structure (upper panel) and changes of number of possible N-glycan conformers at each glycosylated sites (bar graphs) for SlaA with stretched (open) or flapped D5-D6 domains (closed), SlaA Y-body predicted by Alphafold v2.2.0 versus experimental cyroEM, and for monomer in isolation or in the assembled structure. Shown in the bar graphs are changes of glycan conformer numbers at individual glycosylation sites normalized by global changes for all glycans. Positive and negative values indicate an increase or a decrease of possible glycan conformations, respectively, indicative of favourable and unfavuorable entropic contributions. The 7 last N-glycans of the protein were not taken into account for the Alphafold-cryoEM comparison plot, changing the scale of the Y axis compared to the two other plots. Green and red arrows on the right side of the figure indicate a total increase or decrease of glycan conformers during the transition between the two conformations of the protein that were compared in each plot, indicative of favourable and unfavourable entropic contributions to the conformation transition.

**Supplementary figure 9a.**
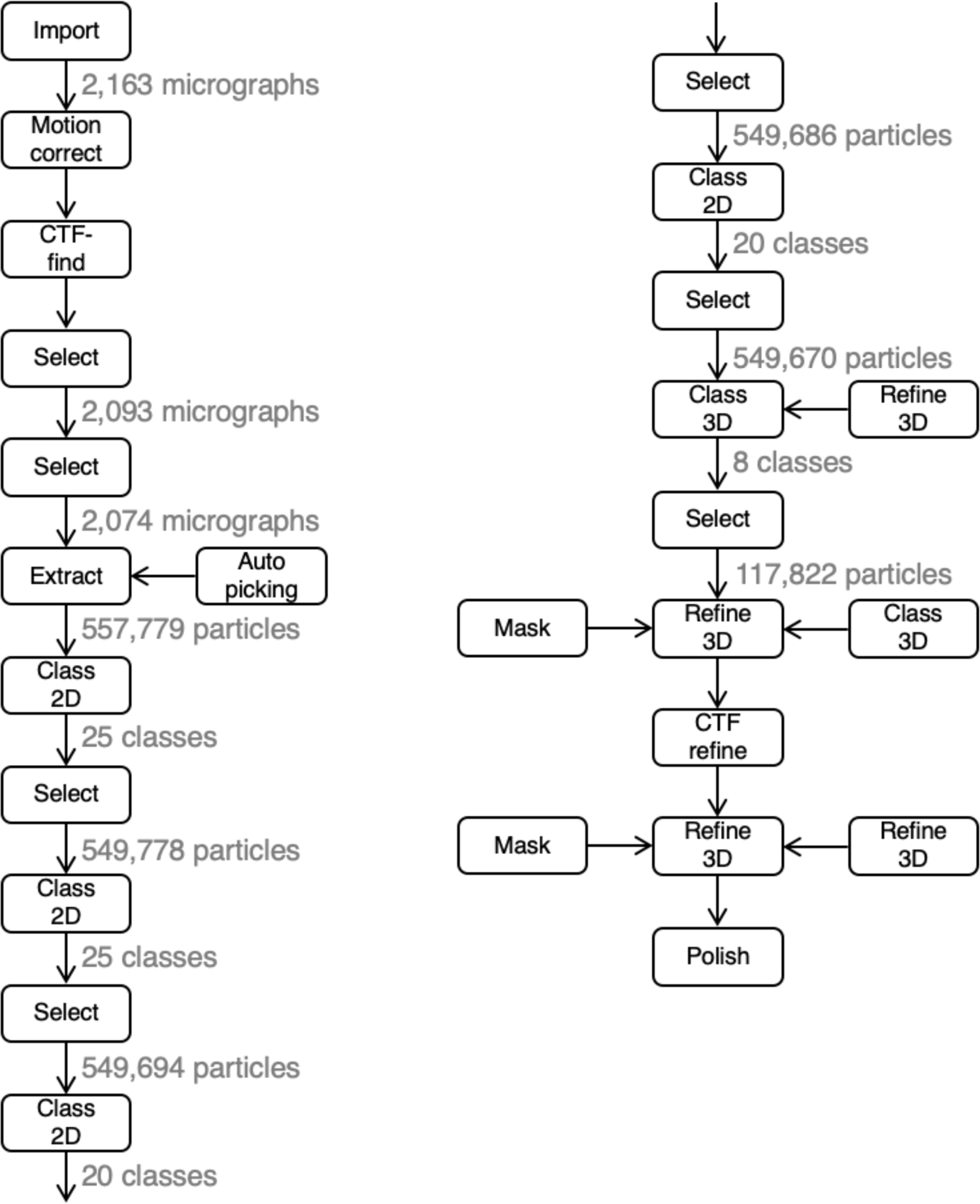
Relion processing workflow for pH 7 dataset (part 1).

**Supplementary figure 9b.**
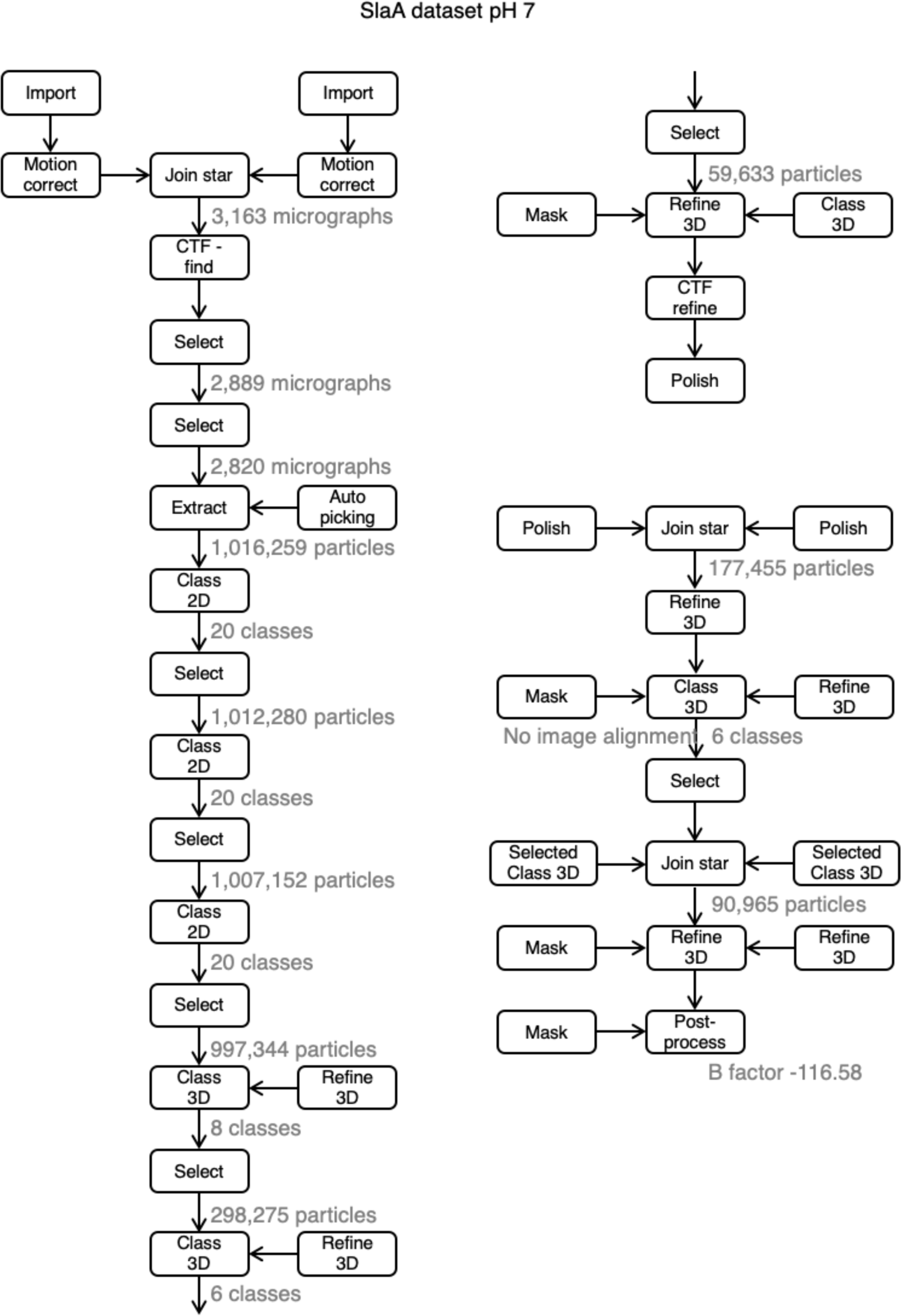
Relion processing workflow for pH 7 dataset (part 2).

**Supplementary figure 10.**
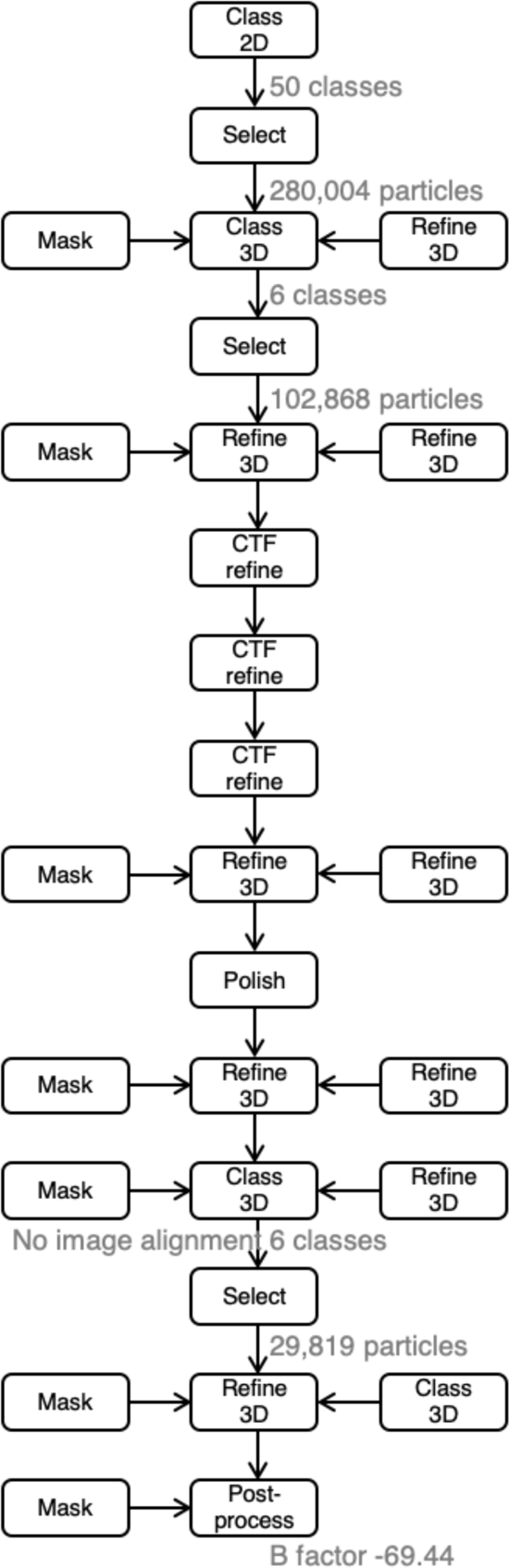
Relion processing workflow for pH 10 dataset.

**Supplementary figure 11.**
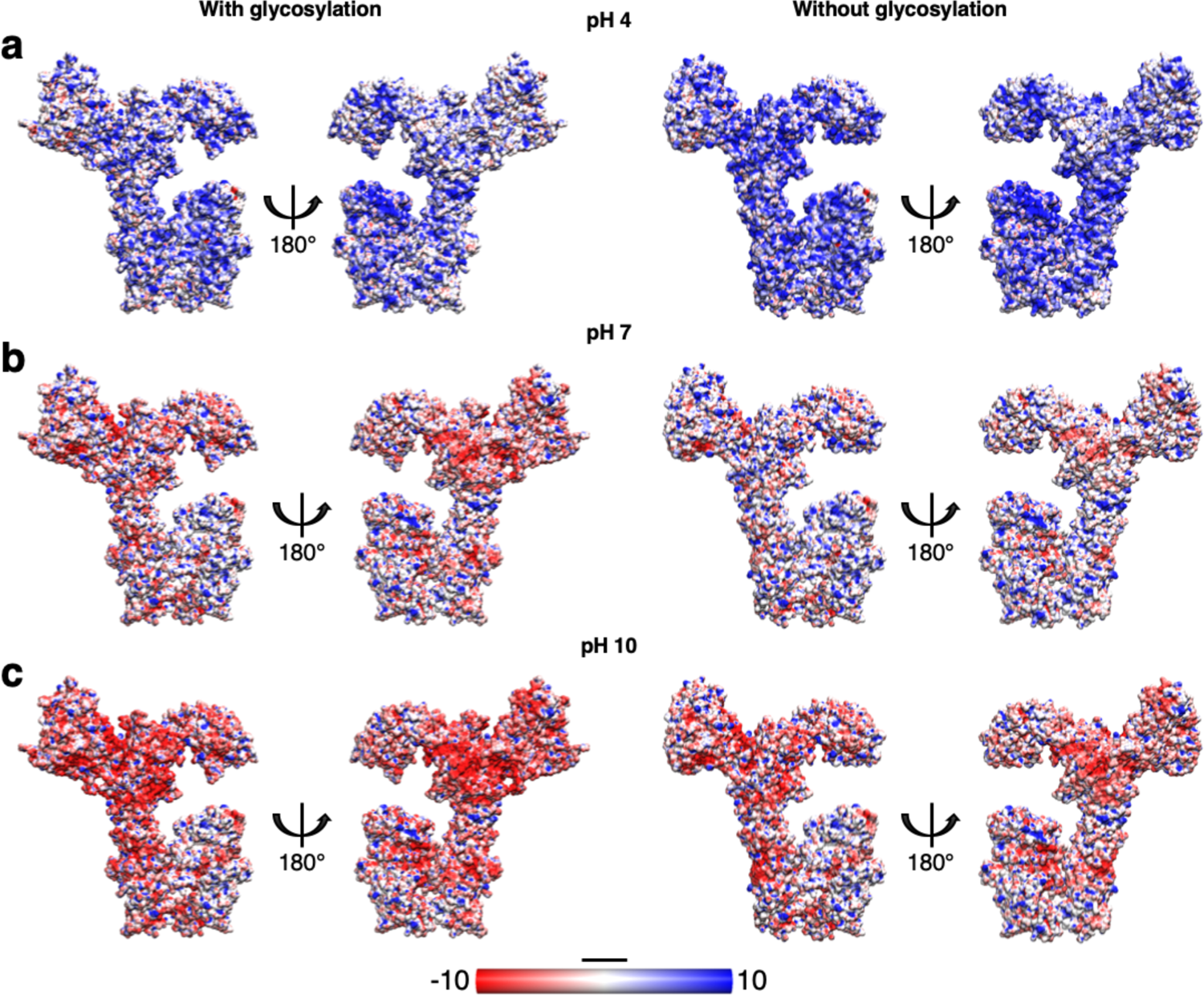
Impact of glycosylation on the electrostatic surface charge of SlaA at different pH values. Comparison of the SlaA electrostatic surface charge with and without glycans at pH 4 (a), 7 (b) and 10 (c). The glycosylation increases the overall surface negative charge of SlaA, particularly noticeable at pH 7 and 10. Scale bar, 20 Å.

**Supplementary figure 12.**
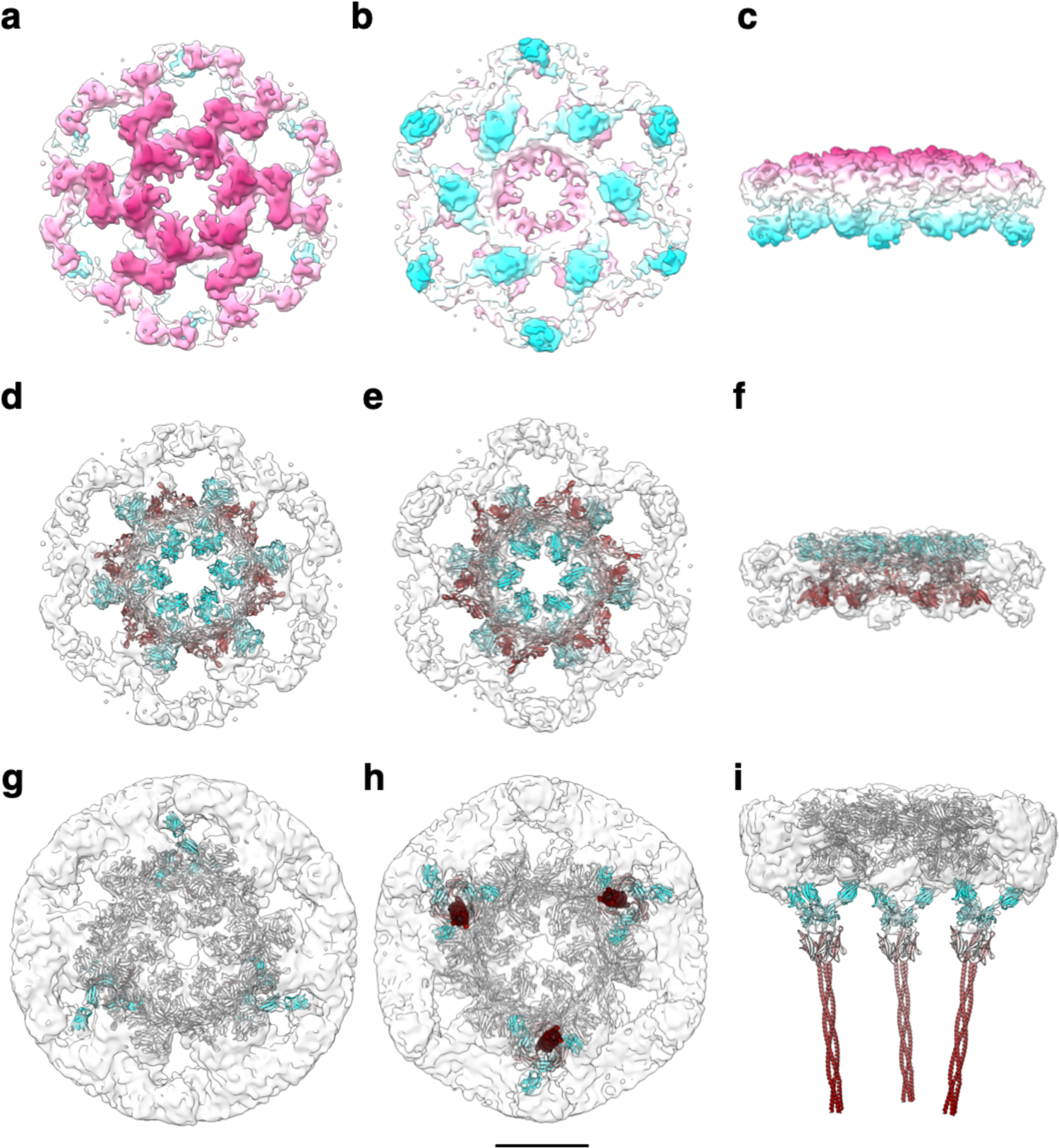
Subtomogram averaging of the S-layer on exosomes. **a-c**, cryoEM density map of the S-layer assembled on exosomes in extracellular (a), intracellular (b), and side (c) views at 8.9 Å resolution. The map shows in magenta the membrane-distal and in cyan the membrane-proximal sides of the lattice. **d-i**, fitting of the SlaA (d-f) and SlaB (g-i) models into the S-layer map in (a-c). SlaA and SlaB are shown in ribbon representation coloured in cyan (N-terminus), grey, and maroon (C-terminus). In (g-i) SlaA is shown in grey. Scale bar, 10 nm.

**Supplementary figure 13.**
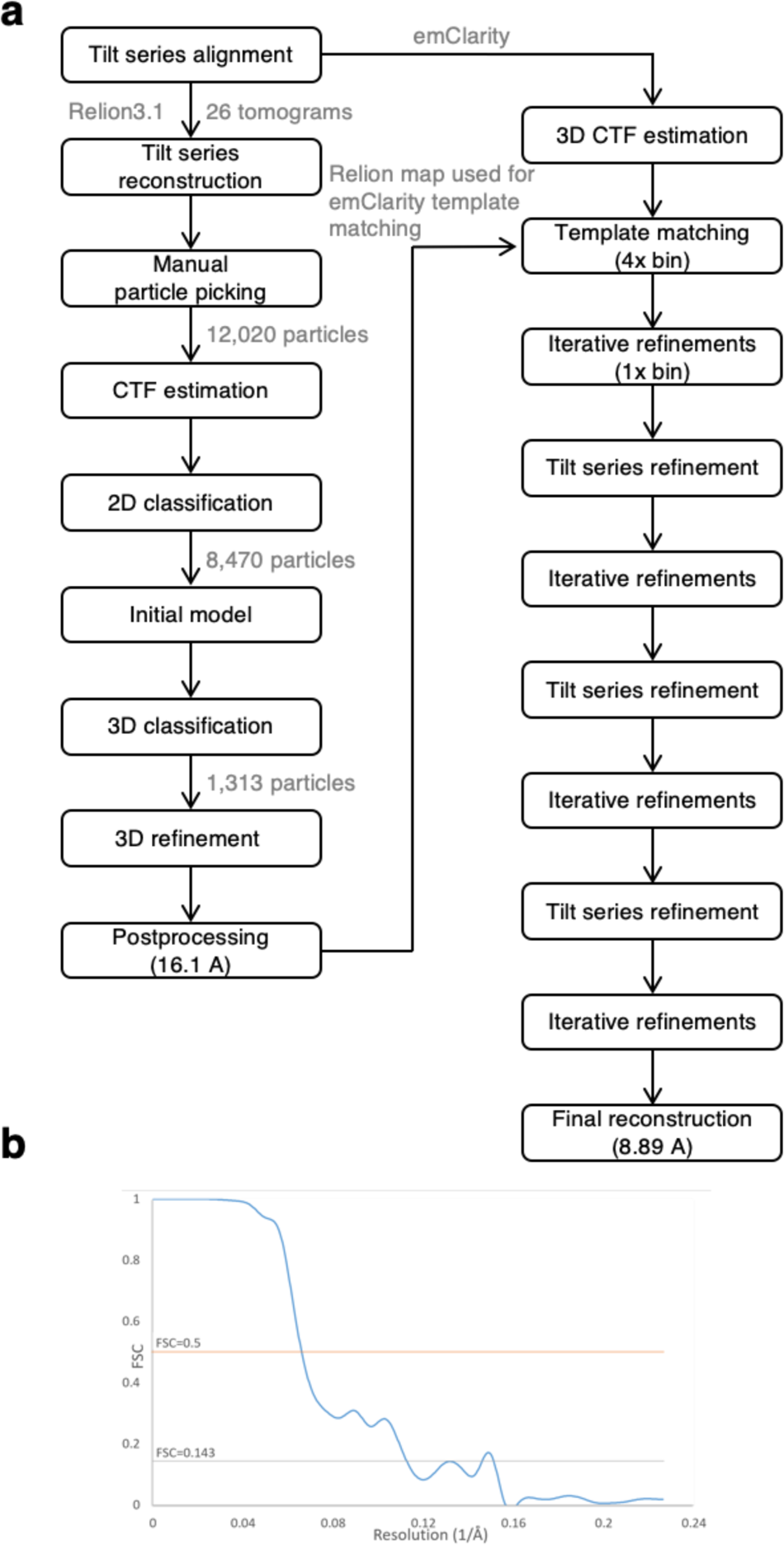
CryoET processing workflow and data quality for cryoEM map of S-layer on exosomes. **a**, subtomogram averaging processing workflow in Relion 3.1 and emClarity. **b**, gold standard FSC of the subtomogram averaging map.

**Supplementary figure 14.**
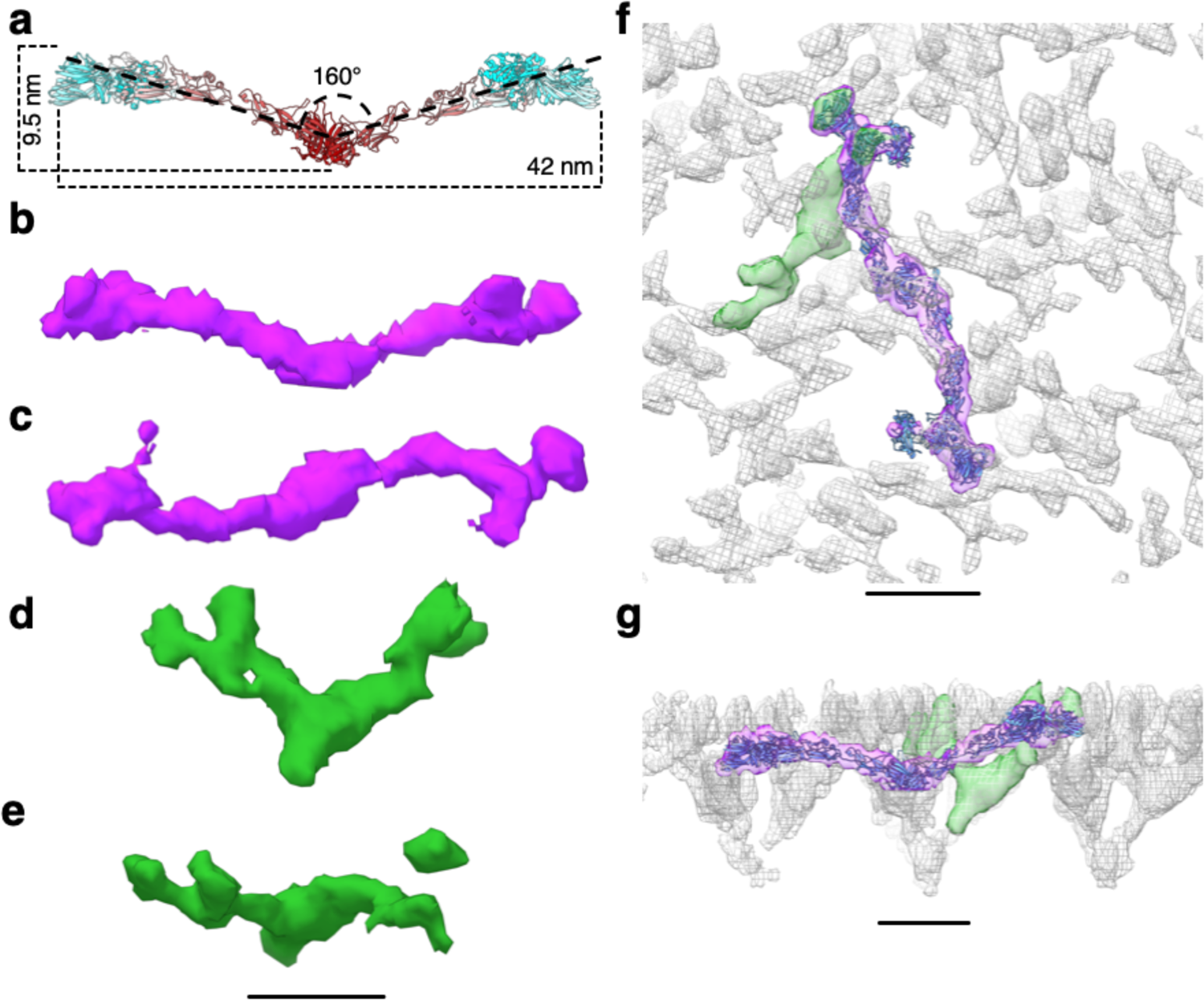
Comparison between current and previously reported^23^ *S. acidocaldarius* SlaA assembly models. **a**, side view of the SlaA dimer (ribbon representation in cyan-grey-maroon from N-terminus to C-terminus). Two SlaA monomers form an angle of 160°. The dimer has a height of 9.5 nm and a length of 42 nm. **b-e**, cryoEM densities extrapolated from the S-layer map published in 2019. The purple density in (b) and (c) (side and extracellular views, respectively) contain the SlaA dimer as presented in this work (f and g). The green density in (d) and (e) (side and extracellular views, respectively), represent the SlaA dimer as reported in 2019. **f** and **g**, show the atomic model of the SlaA dimer in (a) and the cryoEM densities in (b-e) fitting the S-layer cryoEM map (grey mesh) presented in 2019. Scale bar, 10 nm.

**Supplementary figure 15.**
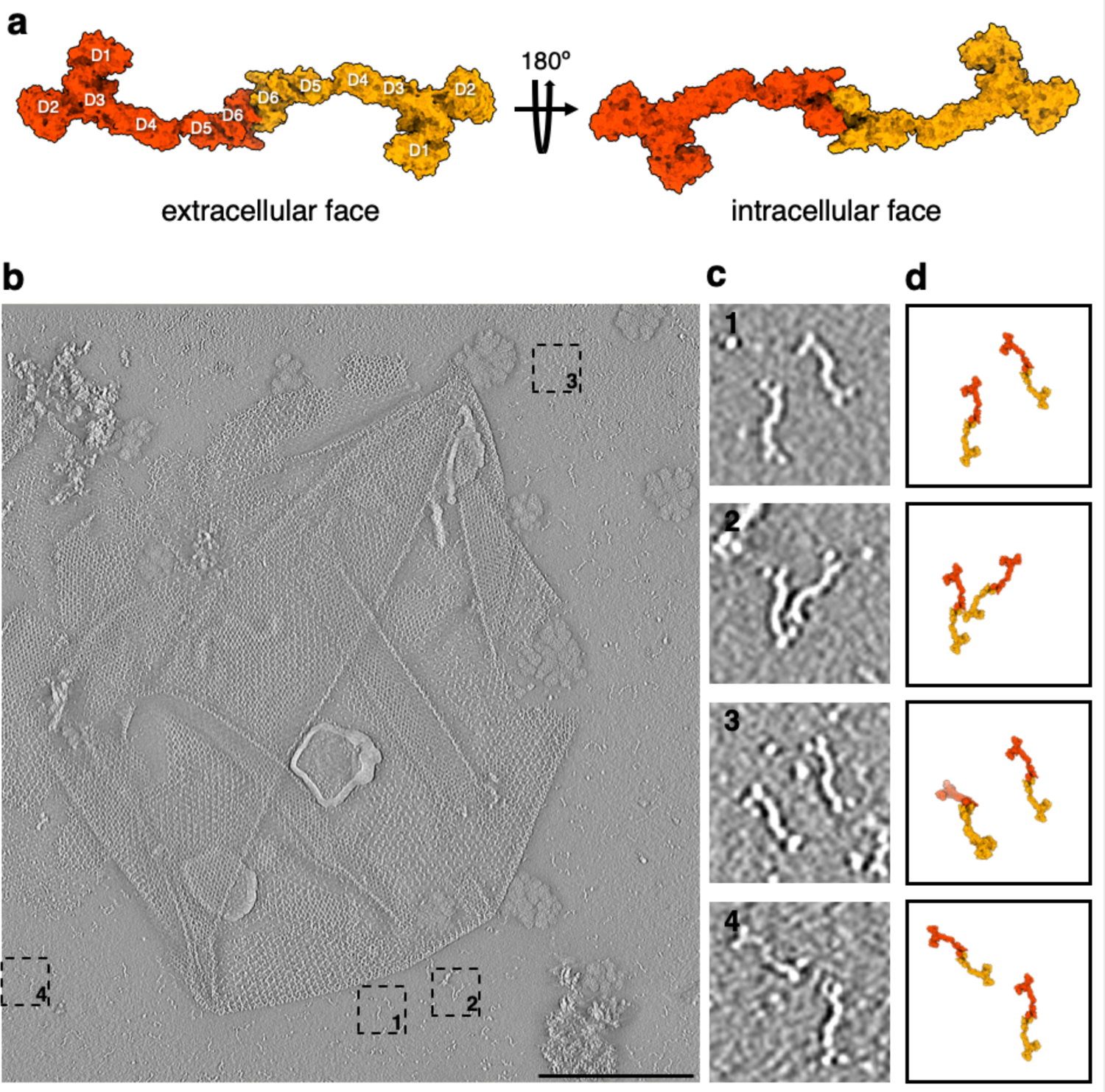
Isolated SlaA-only S-layer from *S. acidocaldarius.* **a**, surface representation of the atomic model of the SlaA dimer, as it occurs in the S-layer. **b**, negative stain electron tomography slice of isolated SlaA-only S-layer. **c**, 1-4 are cut-outs from (b) showing dimeric SlaA with their respective positions marked in (b). **d**, atomic models from (a) scaled and superimposed with the dimers seen in negative stain tomography (c). Scale bar, 200 nm.

**Supplementary figure 16.**
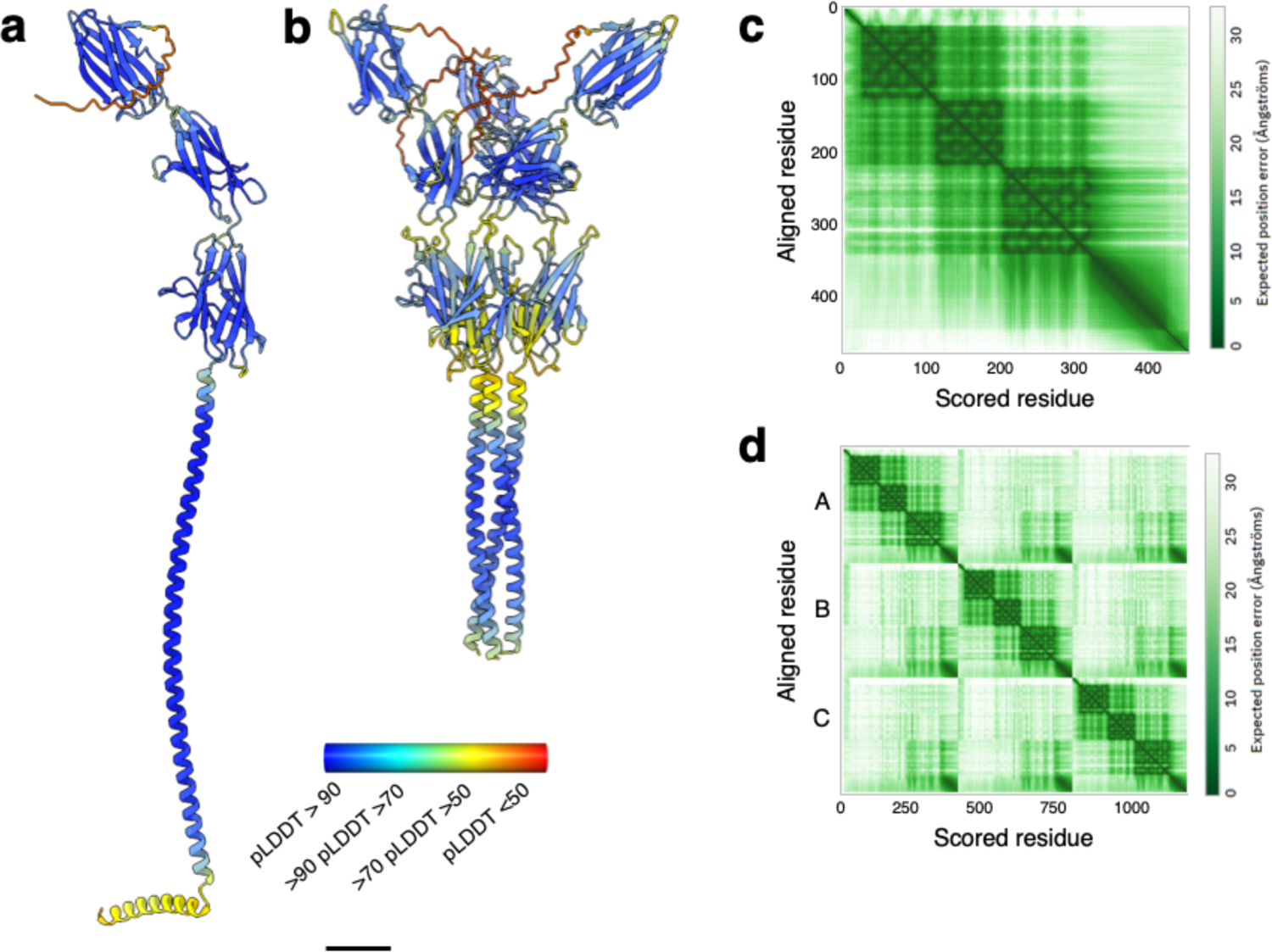
Alphafold v2.2.0 predictions of SlaB monomer and trimer. **a** and **b**, Alphafold v2.2.0 predictions of SlaB monomer and trimer, respectively. The ribbon is coloured by pLDDT (per-residue confidence score) where red indicates very low confidence and blue very high. The trimeric coiled coil of the SlaB trimer (b) is truncated at residue 400, and the complete trimeric coiled coil (Fig. 5d) was predicted using SymmDock. **c** and **d**, PAE (predicted aligned error) plots for (a) and (b), respectively. Scale bar, 20 Å.

**Supplementary figure 17.**
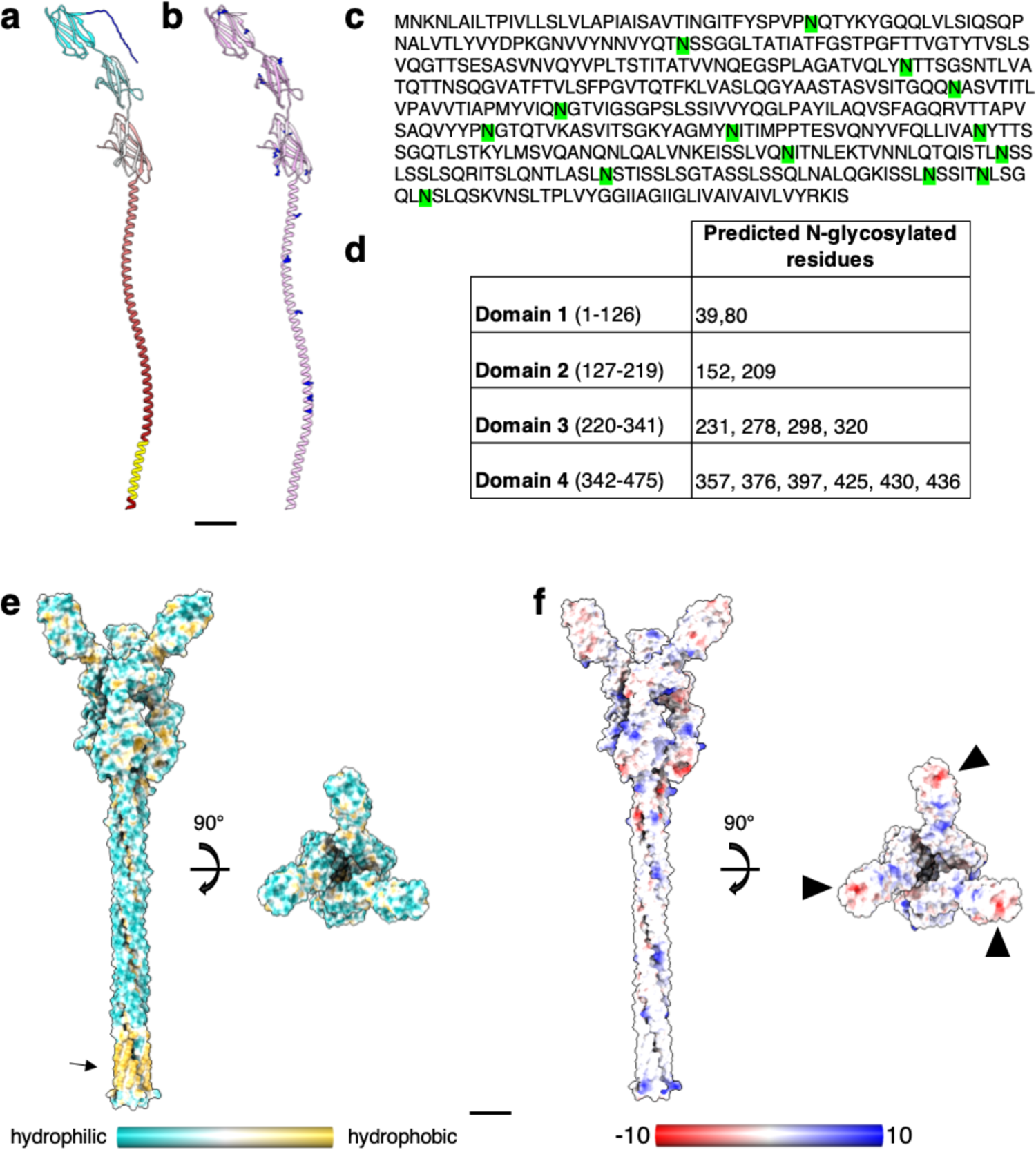
Structural prediction of *S. acidocaldarius* SlaB. **a**, atomic structure of SlaB as predicted by Alphafold v2.2.0 (ribbon representation, cyan-grey-maroon from N-terminus to C-terminus). Amino acids from 1-24 (blue) are predicted as signal peptide by InterPro. Amino acids from 448-470 (yellow) are predicted as trans-membrane helix by TMHMM – 2.0. **b**, putative N-glycosylation sites are labelled as predicted by GlycoPP v1.0. **c**, SlaB sequence with predicted N-glycosylated residues in green. **d**, table showing predicted N-glycosylation distribution across four SlaB domains. **e** and **f**, SlaB trimer (as predicted by Alphafold v2.2.0) surface representation showing hydrophobicity (from most hydrophilic in dark cyan, to white, to most hydrophobic in dark goldenrod) in (e), and electrostatic surface potential (from mostly negative in red, to white, to mostly positive in blue) in (f). The arrow in (e) highlights the predicted hydrophobic trans-membrane region. Arrowheads in f indicate negatively-charged patches that may electrostatically interact with the mostly positively charged SlaA. Scale bar, 20 Å.

**Supplementary figure 18.**
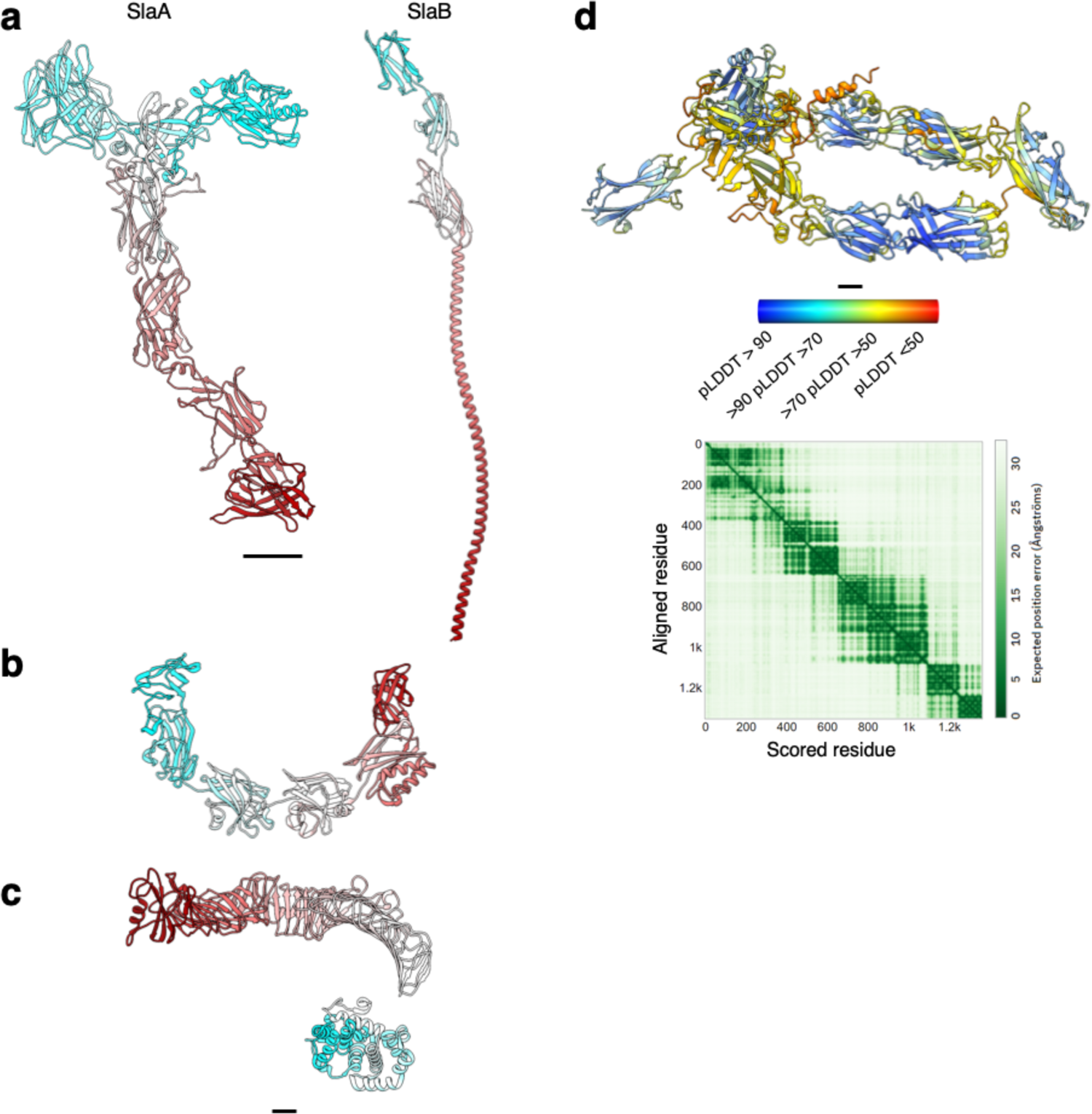
Structure of archaeal and bacterial S-layer proteins. a-c, atomic models are shown in ribbon representation in cyan-grey-maroon from the N-terminus to the C-terminus. *S. acidocaldarius* SlaA (domains D5 and D6 as predicted by Alphafold v 2.2.0) and SlaB (as predicted by Alphafold v.2.2.0) are in (a), *H. volcanii* csg (PDB ID: 7PTR, http://dx.doi.org/10.2210/pdb7ptr/pdb) is in (b), and *C. crescentus* RsaA (N-terminus PDB ID: 6T72, http://dx.doi.org/10.2210/pdb6t72/pdb, C-terminus PDB ID: 5N8P, http://dx.doi.org/10.2210/pdb5n8p/pdb) is in (c). d, SlaA atomic model of *M. sedula* (UNIPROT A4YHQ8, as predicted by Alphafold v2.2.0) and PAE (predicted aligned error) plot. The model is shown in ribbon representation coloured by pLDDT (per-residue confidence score) where red indicates very low confidence and blue very high. Scale bar, 20 Å.

**Supplementary figure 19.**
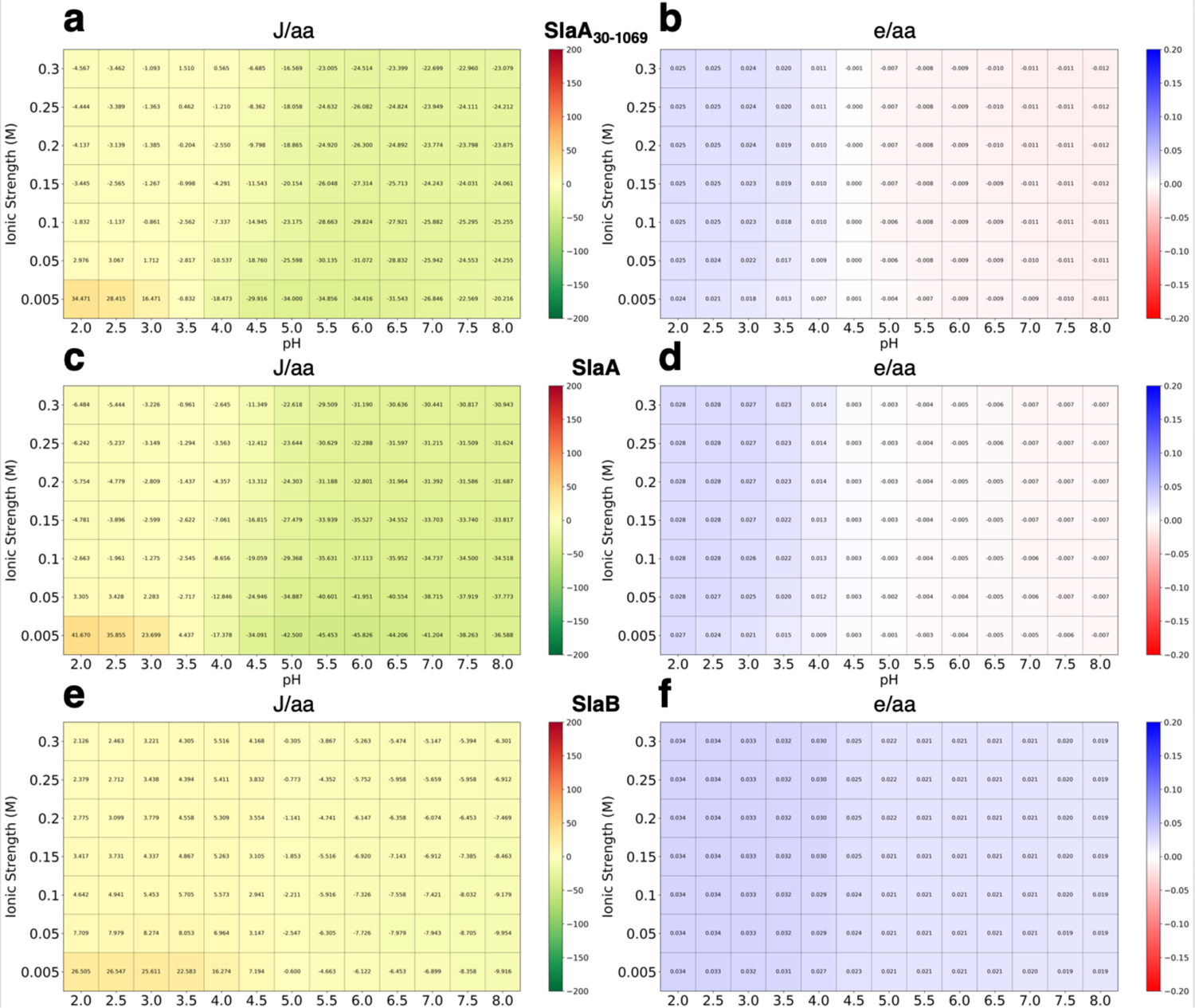
Stability and charge heatmaps for *S. acidocaldarius* SlaA30-1,069, SlaA and SlaB. **a**, **c** and **e**, calculated folded state stability heatmaps for SlaA30-1,069 (a), SlaA (c) and SlaB (e), respectively. SlaA30-1,069, SlaA and SlaB are stable across pH 2-8. **b**, **d** and **f**, calculated charged heatmaps for SlaA30-1,069 (b), SlaA (d) and SlaB (f), respectively. Surface charge shifts from positive to negative for SlaA30-1,069 and SlaA from pH 2 to 8, whereas SlaB shows a largely consistent negative charge.

**Supplementary figure 20.**
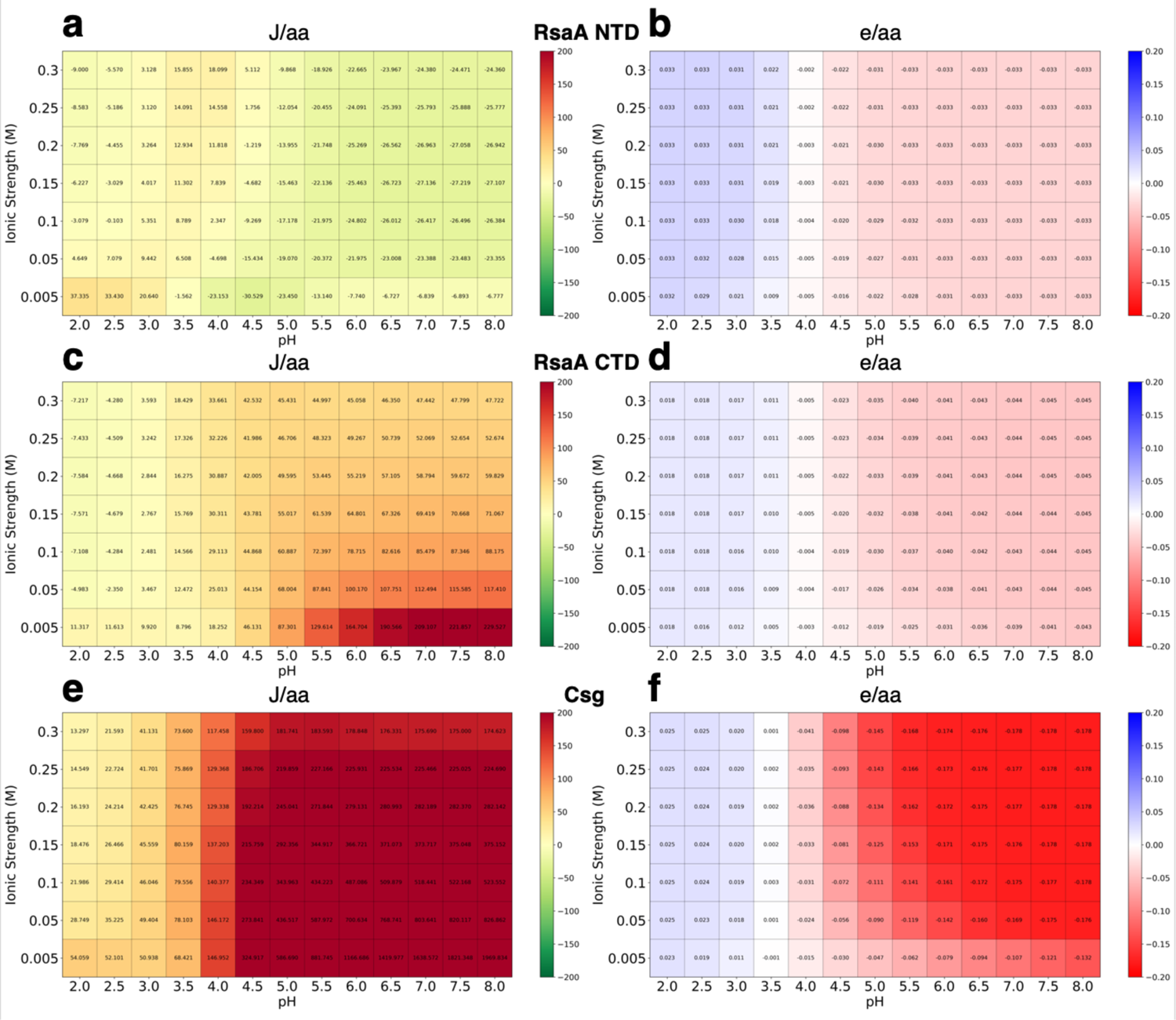
Stability and charge heatmaps for *C. crescentus and H. volcanii* S-layer proteins. **a**, **c** and **e**, calculated folded state stability heatmaps for *C. crescentus* S-layer protein RsaA N-terminus (a) and C-terminus (c) domains, and *H. volcanii* S-layer protein csg (e). The RsaA N-termius domain is largely stable across pH 2-8; RsaA C-terminus domain becomes unstable at elevated pH and low ionic strength; csg’s stability is greatly affected at neutral and high pH. **b**, **d** and **f**, calculated charged heatmaps for RsaA N-terminus (b) and C-terminus (d) domains, and csg (f). RsaA’s and csg’s surface charge shifts from negative to positive from pH 2 to 8. Csg shows a dramatic difference in surface charge from pH 3 to 5.

## Tables

**Supplementary Table 1:**
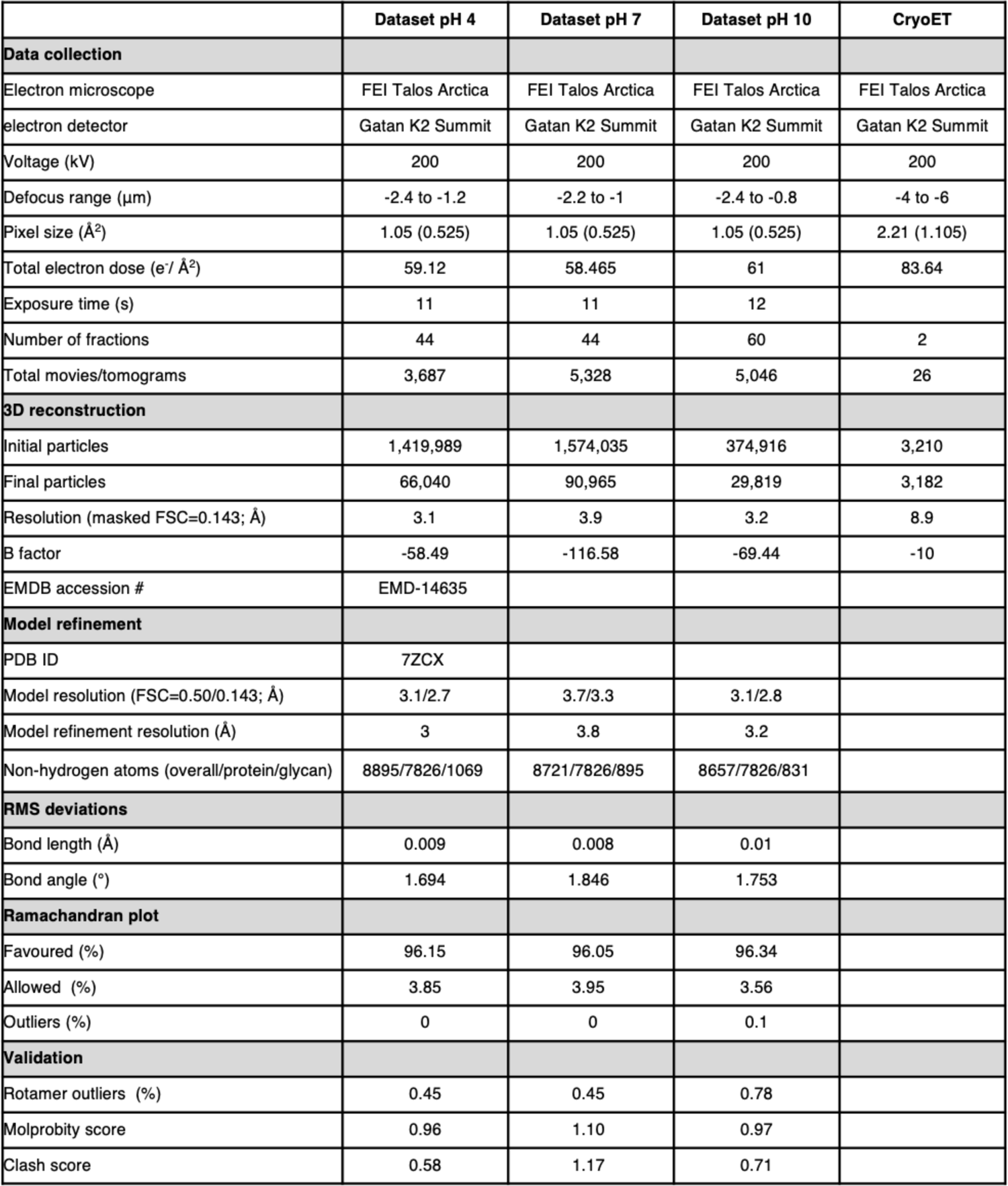
Statistics of data collection, 3D reconstruction and validation.

**Supplementary Table 2:** Mapping of glycan residues from the structure file to residues of the GLYCAM force field or the newly charge-derived SG0 and SG4 residues, representing the 1-substituted and 1,4-substituted SMA.

| Residue | Residue name | Mapped name* | Total charge |
| --- | --- | --- | --- |
| 1201 | NAG | YB4 | 0.000 |
| 1202 | NAG | YBV | -0.194 |
| 1203 | MAN | MB0 | 0.194 |
| 1205 | SMA | SG0 | -0.806 |
| 1301 | NAG | YB4 | 0.000 |
| 1302 | NAG | YB0 | 0.194 |
| 1401 | NAG | YB4 | 0.000 |
| 1402 | NAG | YBQ | -0.388 |
| 1403 | MAN | MB0 | 0.194 |
| 1404 | MAN | MB0 | 0.194 |
| 1405 | SMA | SG4 | -1.000 |
| 1406 | GLC | GB0 | 0.194 |
| 1501 | NAG | YB4 | 0.000 |
| 1502 | NAG | YBQ | -0.388 |
| 1503 | MAN | MB0 | 0.194 |
| 1504 | MAN | MB0 | 0.194 |
| 1505 | SMA | SG0 | -0.806 |
| 1601 | NAG | YB4 | 0.000 |
| 1602 | NAG | YBV | -0.194 |
| 1603 | MAN | MB0 | 0.194 |
| 1605 | SMA | SG0 | -0.806 |
| 1701 | NAG | YB4 | 0.000 |
| 1702 | NAG | YBQ | -0.388 |
| 1703 | MAN | MB0 | 0.194 |
| 1704 | MAN | MB0 | 0.194 |
| 1705 | SMA | SG0 | -0.806 |
| 1801 | NAG | YB4 | 0.000 |
| 1802 | NAG | YBQ | -0.388 |
| 1803 | MAN | MB0 | 0.194 |
| 1804 | MAN | MB0 | 0.194 |
| 1805 | SMA | SG4 | -1.000 |
| 1806 | GLC | GB0 | 0.194 |
| 1901 | NAG | YB4 | 0.000 |
| 1902 | NAG | YBQ | -0.388 |
| 1903 | MAN | MB0 | 0.194 |
| 1904 | MAN | MB0 | 0.194 |
| 1905 | SMA | SG0 | -0.806 |
| 2001 | NAG | YB4 | 0.000 |
| 2002 | NAG | YBQ | -0.388 |
| 2003 | MAN | MB0 | 0.194 |
| 2004 | MAN | MB0 | 0.194 |
| 2005 | SMA | SG0 | -0.806 |
| 2101 | NAG | YB4 | 0.000 |
| 2102 | NAG | YBQ | -0.388 |
| 2103 | MAN | MB0 | 0.194 |
| 2104 | MAN | MB0 | 0.194 |
| 2105 | SMA | SG0 | -0.806 |
| 2201 | NAG | YB4 | 0.000 |
| 2202 | NAG | YB0 | 0.194 |
| 2301 | NAG | YB4 | 0.000 |
| 2302 | NAG | YB6 | 0.000 |
| 2303 | MAN | MB0 | 0.194 |
| 2401 | NAG | YB4 | 0.000 |
| 2402 | NAG | YBQ | -0.388 |
| 2403 | MAN | MB0 | 0.194 |
| 2404 | MAN | MB0 | 0.194 |
| 2405 | SMA | SG4 | -1.000 |
| 2406 | GLC | GB0 | 0.194 |
| 2501 | NAG | YB4 | 0.000 |
| 2502 | NAG | YBQ | -0.388 |
| 2503 | MAN | MB0 | 0.194 |
| 2504 | MAN | MB0 | 0.194 |
| 2505 | SMA | SG0 | -0.806 |
| 2601 | NAG | YB4 | 0.000 |
| 2602 | NAG | YBV | -0.194 |
| 2603 | MAN | MB0 | 0.194 |
| 2605 | SMA | SG0 | -0.806 |
| 2701 | NAG | YB4 | 0.000 |
| 2702 | NAG | YBQ | -0.388 |
| 2703 | MAN | MB0 | 0.194 |
| 2704 | MAN | MB0 | 0.194 |
| 2705 | SMA | SG0 | -0.806 |
| 2801 | NAG | YB0 | 0.194 |
| 2901 | NAG | YB4 | 0.000 |
| 2902 | NAG | YBQ | -0.388 |
| 2903 | MAN | MB0 | 0.194 |
| 2904 | MAN | MB0 | 0.194 |
| 2905 | SMA | SG0 | -0.806 |
| 3001 | NAG | YB4 | 0.000 |
| 3002 | NAG | YBQ | -0.388 |
| 3003 | MAN | MB0 | 0.194 |
| 3004 | MAN | MB0 | 0.194 |
| 3005 | SMA | SG0 | -0.806 |

**Supplementary Table 3:** RESP charges derived for residue SG0 on the HF/6-31G*//HF/6-31G* level of theory (see Methods for details).

| Atom | Charge |
| --- | --- |
| C6 | -0.0742 |
| S1 | 1.2319 |
| O6 | -0.7039 |
| O7 | -0.7039 |
| O8 | -0.7039 |
| C5 | 0.1334 |
| O5 | -0.2985 |
| C1 | 0.3664 |
| C2 | 0.0949 |
| O2 | -0.2312 |
| C3 | 0.324 |
| O3 | -0.2189 |
| C4 | 0.1802 |
| O4 | -0.2023 |

**Supplementary Table 4:** RESP charges derived for residue SG4 on the HF/6-31G*//HF/6-31G* level of theory (see Methods for details).

| Atom | Charge |
| --- | --- |
| C6 | -0.0559 |
| S1 | 1.1863 |
| O6 | -0.6936 |
| O7 | -0.6936 |
| O8 | -0.6936 |
| C5 | 0.1413 |
| O5 | -0.2759 |
| C8 | 0.3583 |
| C2 | 0.1387 |
| O2 | -0.2298 |
| C3 | 0.2118 |
| O3 | -0.1919 |
| C4 | 0.1517 |
| O4 | -0.3538 |

## Supplementary Movies

**Supplementary Movie 1.** Atomic structure and glycosylation of SlaA_30-1069_. The SlaA_30-1069_ cryoEM map is shown in cornflower blue. The atomic structure is shown in ribbon representation in cyan-grey-maroon from the N-terminus to the C-terminus. The glycosylated Asn residues are in orange and the glycans are represented as ball-stick in steel blue, with N atoms in blue, O in red and S in yellow.

**Supplementary Movie 2.** Flexibility of the D1, D5 and D6 domains. Sequence of 2D classifications obtained in Relion 3 of negatively stained SlaA. D1-4 were aligned, showing the flexibility of D1, D5 and D6.

**Supplementary Movie 3.** Comparison of SlaA_30-1069_ structure at pH 4 and 10. r.m.s.d. alignment between SlaA_30-1069_ atomic models at pH 4 and pH 10. Smaller deviations are shown in blue and larger deviations in red, with mean r.m.s.d. = 0.79 Å, as in Figure 3b.

**Supplementary Movie 4.** Model of the assembled *S. acidocaldarius* S-layer. The SlaA models are shown in blue-grey-purple ribbons and the SlaB models are in red-orange-yellow ribbons.

